# Unsaturated fatty acids augment protein transport via the SecA:SecYEG translocon

**DOI:** 10.1101/2021.03.26.437172

**Authors:** Michael Kamel, Maryna Löwe, Stephan Schott-Verdugo, Holger Gohlke, Alexej Kedrov

## Abstract

The translocon SecYEG forms the primary protein-conducting channel in the cytoplasmic membrane of bacteria, and the associated ATPase SecA provides the energy for the transport of secretory and cell envelope protein precursors. The translocation requires negative charge at the lipid membrane surface, but its dependence on the properties of the membrane hydrophobic core is not known. Here, we demonstrate that SecA:SecYEG-mediated protein transport is immensely stimulated by unsaturated fatty acids (UFAs). Furthermore, UFA-rich tetraoleoyl-cardiolipin, but not bis(palmitoyloleoyl)-cardiolipin, facilitate the translocation via the monomeric translocon. Biophysical analysis and molecular dynamics simulations show that UFAs determine the loosely packed membrane interface, where the N-terminal amphipathic helix of SecA docks. While UFAs do not affect the translocon folding, they promote SecA binding to the membrane, and the effect is enhanced manifold at elevated ionic strength. Tight SecA:lipid interactions convert into the augmented translocation. As bacterial cells actively change their membrane composition in response to their habitat, the modulation of SecA:SecYEG activity via the fatty acids may be crucial for protein secretion over a broad range of environmental conditions.

## Introduction

Protein transport across the cytoplasmic bacterial membrane is an essential step in biogenesis of cell envelope and secretory proteins [1]. Most of these proteins cross the membrane post-translationally as unfolded precursors (preproteins). The preproteins bear a cleavable N-terminal hydrophobic signal sequence, which slows down their folding in the cytoplasm. The preproteins are picked up by holdase chaperones, such as SecB, and delivered in the largely unfolded state to the Sec machinery (Figure 1A). The core of the Sec machinery consists of the heterotrimeric membrane-embedded channel, or *translocon*, SecYEG and the membrane-associated ATPase SecA. The translocon builds a narrow transmembrane conduit for the unfolded preproteins, and it is primed by insertion of the signal sequence at the translocon:lipid interface. The activity of the SecYEG-bound ATPase SecA provides the energy for directional transport of the preprotein through the translocon [2].

**Figure 1.**
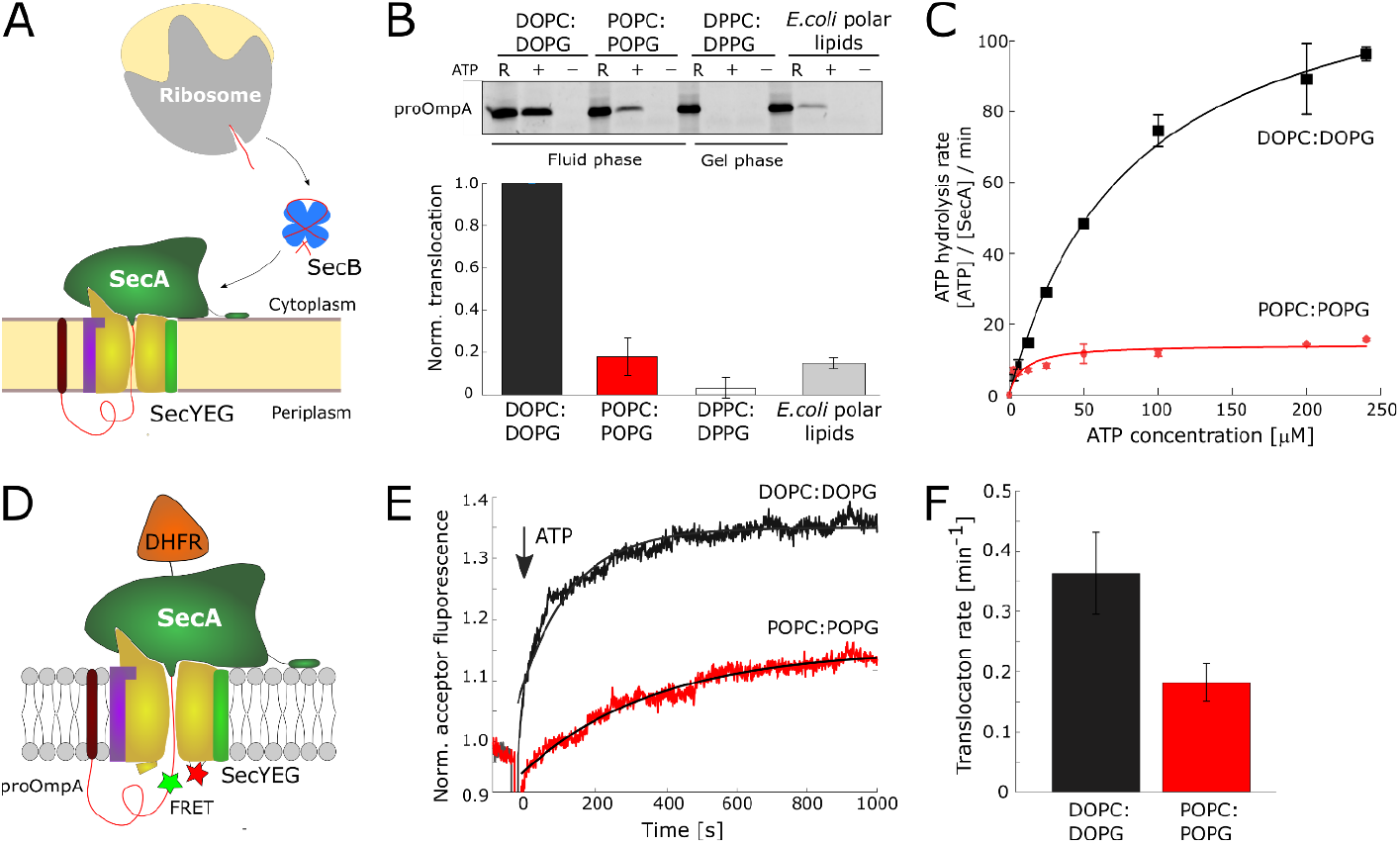
Unsaturated fatty acids stimulate Sec-mediated translocation. **(A)** Primary pathway for protein translocation is composed of the cytoplasmic holdase chaperone SecB, the membrane-associated ATPase SecA, and the transmembrane translocon SecYEG. **(B)** Translocation of the preprotein proOmpA is sensitive to the content of the unsaturated fatty acids (UFAs) in the membrane core. The translocation efficiency in DOPC:DOPG bilayers was used for normalization. Error bars show standard deviation (SD) values, as measured in triplicates. **(C)** The ATPase activity of SecA associated with proOmpA translocation is strongly stimulated in DOPC:DOPG proteoliposomes (K_M_ of 83 ± 6 μM). **(D)** Scheme of the stable translocation intermediate. Unfolded proOmpA domain is translocated via SecA:SecYEG, but the folded domain DHFR stalls the transport and jams the translocon. Two fluorophores positioned within proOmpA and at the periplasmic side of SecY allow for FRET once the stalled complex is formed. **(E)** Assembly of the translocation intermediate is followed as the fluorescence of the FRET acceptor is increasing with time after addition of ATP (arrow). The UFA-enriched DOPC:DOPG membranes (black) stimulate the proOmpA translocation and ensure formation of the stalled translocation intermediate. **(F)** Apparent translocation rates determined from the FRET-based assay validate the faster translocation reaction in the UFA-enriched DOPC:DOPG membrane.

Being an essential and universally conserved system, the Sec machinery has been extensively investigated, and the importance of the lipid environment has been highlighted for posttranslational translocation and interactions of the translocon with ribosomes [3–6]. Anionic lipids, mainly phosphatidylglycerol (PG) and cardiolipin (CL), appear to be essential for the preprotein secretion in bacteria, as they form electrostatic interactions with SecA, enabling the formation of the active SecA:SecYEG complex. The N-terminus of SecA forms an amphipathic helix, whose basic residues interact with the anionic lipids and anchor the ATPase to the membrane, even in the absence of SecYEG [3,7]. While various anionic lipids may support the translocation [7], the requirement for particular species, such as CL, has remained intensively disputed [7–9].

While the role of lipid head groups for the dynamics of the SecA:SecYEG machinery has been thoroughly examined, only limited insights on the effect of the constituting fatty acids are available. Initial reconstitution of SecYEG in proteoliposomes revealed that dioleoyl-phosphatidylethanolamine (DOPE) stimulates preprotein translocation [5]. The effect was attributed to the conical shape of DOPE molecules built of two mono-unsaturated fatty acids (UFAs) and the small head group, but the mechanistic explanation of the stimulation has been lacking. This limited knowledge contrasts the essential complexity of cellular membranes, where the diversity of fatty acids arises from variations in their length and the unsaturation level. Furthermore, cells regulate the UFA content in response to changing environmental factors, the habitat style, and the growth phase, while the lipid head group distribution remains rather constant [10]. As an example, the elevated UFA content allows to maintain the fluidity of cellular membranes at low temperatures, so the ratio of UFAs to saturated fatty acids in the inner membrane of *E. coli* changes from 1:1 to 2:1 when the growth temperature is reduced to 17 °C [11].

Several membrane-associated protein complexes in bacteria and eukaryotes appear to be sensitive to the UFA content [12–14]. The sensitivity is mediated either by their membrane-embedded subunits or via altered interactions at the membrane:protein interface, and it may facilitates UFA-specific signalling reactions, protein folding and degradation. Here, we demonstrate for the first time that changes in the UFA content have an immense effect on SecA:SecYEG-mediated protein translocation. Increasing the *cis*-UFA content within the fluid phase membrane leads to a manifold increase in the translocon activity. Biophysical analysis and all-atom molecular dynamics simulations show that the structure of the fatty acids does not affect SecYEG stability, but UFAs determine a loosely packed membrane interface and facilitate apolar SecA:lipid interactions. The stimulated association of the ATPase with the lipid membrane leads to the augmented activity of the SecA:SecYEG complex. We further demonstrate that the UFA-enriched tetraoleoyl-CL also stimulates the translocation, and the stimulation does not involve oligomerization of SecYEG. Our results reveal that the organization of the lipid membrane plays a prominent role in the regulation of protein translocation and suggest that the regulation may be employed upon the bacterial adaptation to various habitat conditions.

## Results

### Unsaturated fatty acids stimulate protein translocation

The effect of the fatty acid structure on SecA:SecYEG-mediated translocation was examined using liposomes of defined and tailored lipid compositions. The inner membranes of *E. coli* are composed mainly of the zwitterionic lipid phosphatidylethanolamine (PE; up to 70 mol %) and anionic lipids, PG (20-25 mol%) and CL (3-5 mol %) [15]. Due to the small radius of their head groups, some PE species, such as DOPE, do not form planar lipid bilayers *in vitro*, and their gel-to-fluid transition temperatures are substantially higher than those of PG or another zwitterionic lipid phosphatidylcholine (PC) with the same fatty acid composition. Thus, to avoid non-lamellar structures and phase separation in the composite liposomes, PC lipids were employed here as the zwitterionic component. SecYEG was reconstituted into liposomes containing 30 mol % PG and 70 mol % PC; that way, the fraction of anionic lipids (PG) mirrored the abundance in the inner membrane of *E. coli* and should be sufficient to enable the electrostatically driven SecA:lipid interactions [3,5]. Indeed, the translocon remained active when reconstituted into POPC:POPG lipids, thus validating PC:PG liposomes as a functional membrane mimetic (Suppl. Figure 1A) [5].

To examine the effect of UFAs on translocation, the fatty acid composition of the proteoliposomes was varied, while keeping the PC:PG molar ratio of 7:3 constant. Phospholipids composed of dipalmitoyl (16:0/16:0, DP; no mono-UFA), 1-palmitoyl-2-oleoyl (16:0/18:1 Δ9-*cis*, PO; 50 % mono-UFA), and dioleoyl (18:1/18:1 Δ9-*cis*, DO; 100 % mono-UFA) fatty acids, as well as the natural extract of *E. coli* polar lipids (average mono-UFA content ~50 mol %), were tested (Figure 1B). Fully saturated fatty acids of DPPC:DPPG lipids resulted in a gel-phase membrane (transition temperatures 42 °C), but both POPC:POPG and DOPC:DOPG membranes (phase transition temperatures −2 °C and −18 °C, respectively) were present in the disordered fluid phase, as confirmed by the fluorescence anisotropy analysis of the conventional bilayer-incorporated reporter DPH (Suppl. Figure 1B) [16]. The membrane fluidity was essential for SecA:SecYEG-mediated translocation at 37 °C: Transport of the preprotein proOmpA was nearly zero in the gel-phase DPPC:DPPG liposomes; hence, the essential SecA:SecYEG dynamics must be severely suppressed. The fluid-phase membranes ensured the preprotein translocation, but the translocation efficiency manifested striking variations: DOPC:DOPG lipids strongly stimulated the activity of SecA:SecYEG, as up to 5-fold more proOmpA was accumulated in liposomes, in comparison to POPC:POPG membranes (Figure 1B). The translocation efficiency correlated with the ATPase activity of SecA (Figure 1C), suggesting that the length of the polypeptide chain translocated per cycle of ATP hydrolysis was not altered. The translocation in native *E. coli* extracts enriched with partially unsaturated PE lipids was substantially lower than in DOPC:DOPG liposomes, so that the stimulatory effect must originate from the fatty acid structure and presence of UFAs, but not the head group composition.

To investigate whether UFAs modulate the translocation rate, we analysed the kinetics of a single translocation cycle using the Förster’s resonance energy transfer (FRET)-based transport assay [9]. The N-terminal proOmpA domain of the fusion preprotein proOmpA-DHFR is translocated into the liposome lumen (Figure 1D). The preprotein stays trapped within the Sec complex, once the folded C-terminal DHFR domain blocks the SecA:SecYEG machinery. The single translocation cycle results in a stable translocation intermediate. Once the translocation intermediate is assembled, the fluorophores in the C-terminal part of proOmpA (Cyanine3, donor) and at the periplasmic side of SecYEG (Atto 643, acceptor) come into proximity allowing for FRET. The increase in the acceptor fluorescence is assigned to the intermediate assembly and, once recorded over time, it provides an insight into the translocation kinetics [9]. SecYEG reconstituted in the UFA-enriched DOPC:DOPG liposomes displayed approx. twofold higher rates of the intermediate formation compared to SecYEG in POPC:POPG liposomes (Figure 1E and 1F). The lower amplitude of the FRET signal observed in POPC:POPG further suggested that a fraction of SecA:SecYEG complexes did not completely translocate the proOmpA domain, likely due to the slower translocation kinetics accompanied by inactivation of the temperature-labile SecA ATPase [17]. Together, the results of the functional assays reveal that UFAs within the physiologically fluid lipid membrane stimulate the efficiency of the SecA:SecYEG translocon and increase the rate of the polypeptide chain transport.

### SecYEG stability and topology are not affected by fatty acid composition

The prominent effect of the fatty acid composition on Sec-mediated translocation suggests that the hydrophobic core of the membrane could either affect stability and dynamics of the SecYEG translocon or be a novel factor that regulates SecA binding and SecA:SecYEG assembly. To probe the translocon stability in various environments, we established differential scanning fluorimetry (DSF) measurements, which report on the protein denaturation based on changes in the fluorescence emission of tryptophan residues [18]. Loss of the native protein structure leads to exposure of tryptophan residues to the aqueous solvent, so that their fluorescence is red-shifted. The SecYEG translocon contains eight tryptophan residues positioned at the ends of transmembrane helices (Suppl. Figure 2A), and their fluorescence was recorded over the temperature range from 25 to 80 °C. An abrupt change observed both in detergent micelles and in liposomes indicated the cooperative denaturation of the translocon (Figure 2A). Notably, the lipid environment greatly stabilized SecYEG: The denaturation temperature *T*_m_ in DDM micelles was measured at 47 °C, but it increased to 67 °C in DOPC:DOPG liposomes (Figure 2B). Variations in the UFA content had a minor effect on the *T*_m_ value (Figure 2B) indicating that SecYEG was equally stable and correctly folded in the examined lipid bilayers. As the lipid composition did not affect the reconstitution efficiency of the translocon, and POPC:POPG lipids rather favoured its functional topology in lipid membranes (Suppl. Figure 2B and C), UFA-specific SecYEG inactivation upon the reconstitution was excluded. Importantly, a recent mass-spectrometry analysis of SecYEG-associated fatty acids in native membranes did not reveal deviations from the overall UFA distribution in *E. coli* inner membranes, so that the translocon does not form preferential interactions with specific fatty acids [19]. Finally, the SecYEG^prlA4^ mutant, which demonstrates elevated preprotein translocation due to the altered structure of the central pore [20], was similarly sensitive to the UFA content as the wild-type translocon (SecYEG^WT^, Figure 2C). Thus, the effect of UFAs on SecA:SecYEG-mediated translocation was dominant and not related to SecYEG:lipid contacts, but possibly originated from altered protein:lipid interactions at the membrane interface.

**Figure 2.**
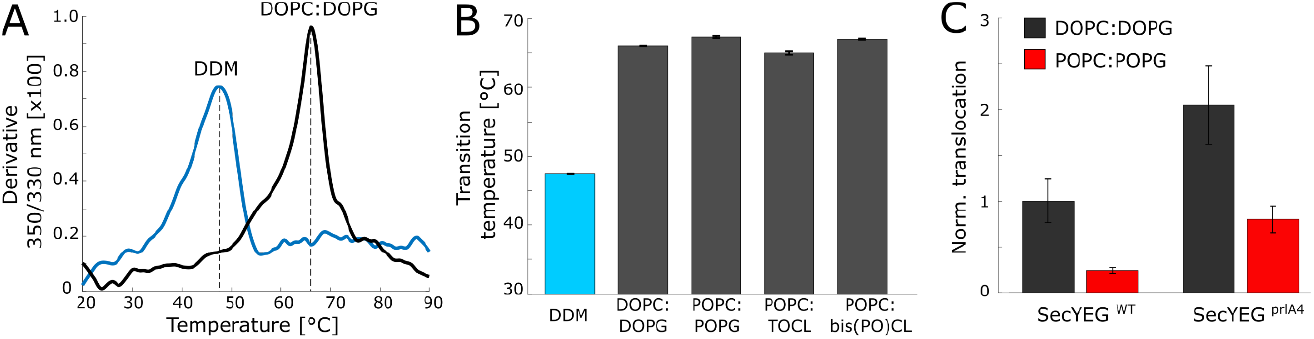
The fatty acids content does not affect the stability and dynamics of the SecYEG translocon. **(A)** Profiles of the thermal denaturation of SecYEG, as measured by differential scanning fluorometry based on the changes in the intrinsic tryptophan fluorescence. The lipid membrane substantially stabilizes the reconstituted translocon. **(B)** Thermal stability of SecYEG in liposomes is not influenced by the fatty acid composition of the membrane. Error bars show SD values, as measured in duplicates. **(C)** The activity of the up-regulated translocon mutant SecYEG^prlA4^ is equally sensitive to the UFA content as the wild-type translocon (SecYEG^WT^), suggesting that UFAs play a SecYEG-independent role in the translocation. Error bars show SD values, as measured in duplicates.

### Unsaturated fatty acids cause loose packing of lipid head groups

To study whether UFAs alter the lipid membrane organization at the interface, we employed the environment-sensitive dye laurdan. The dye spontaneously intercalates between the lipid head groups, and its general fluorescence polarization value decreases with higher water permeation, which is characteristic for loose lipid packing [21]. The fluorescence polarization measured in DOPC:DOPG liposomes was significantly lower than that in POPC:POPG membranes (−0.52±0.02 vs. −0.40±0.01; Figure 3A), thus suggesting more disordered interface structure for the UFA-enriched lipid bilayer.

**Figure 3.**
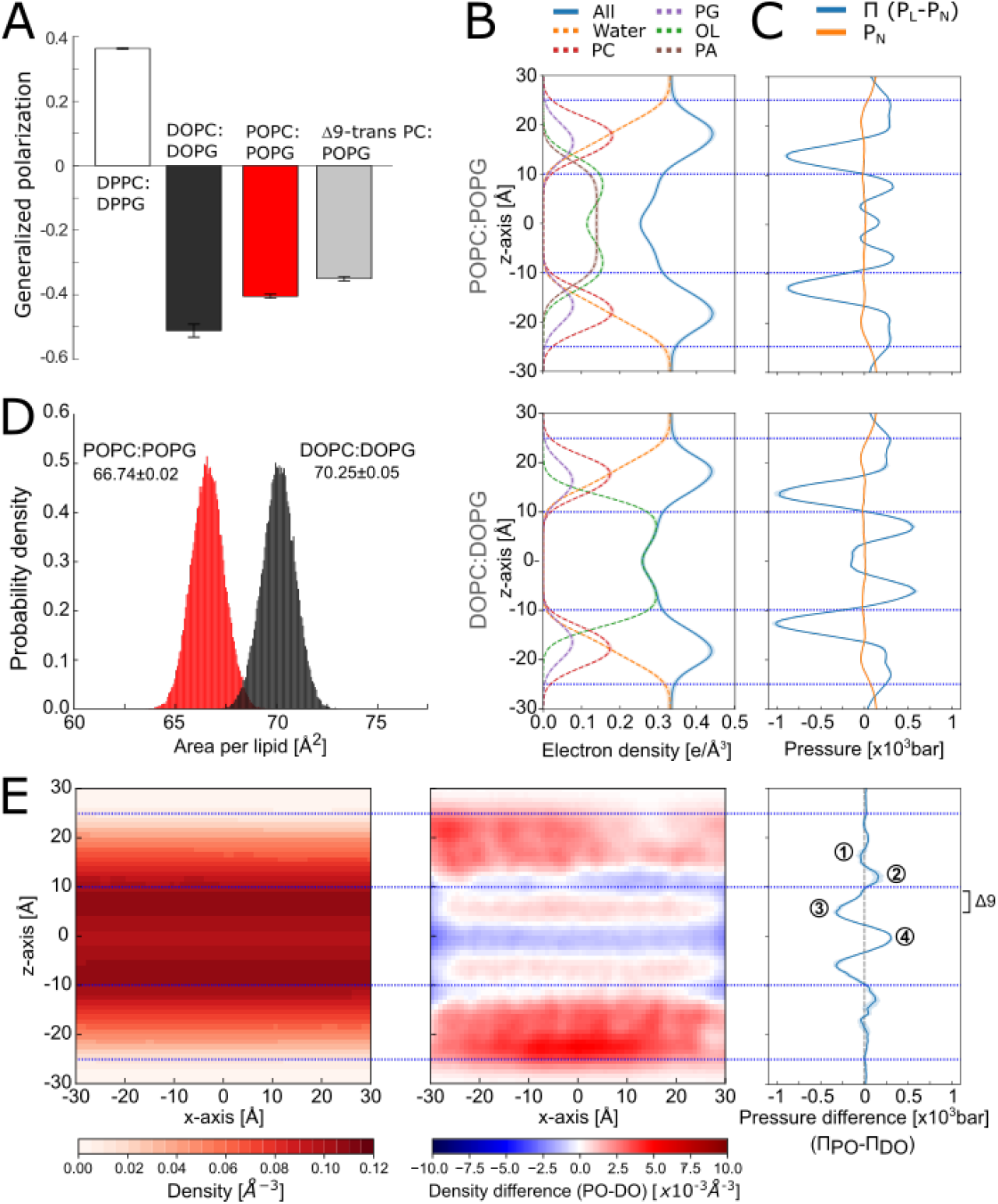
Fatty acid structure determines the biophysical properties of lipid bilayers. **(A)** General polarization of the laurdan dye suggests a looser packing of the lipid head groups in *cis*-UFA-enriched lipid bilayers (DOPC:DOPG). Error bars show SD values, as measured in triplicates. **(B)** Electron density profiles of POPC:POPG (top) and DOPC:DOPG (bottom) bilayers. The profiles provide information regarding the location of the membrane components along the plane normal. PC: phosphatidylcholine headgroup; PG: phosphatidylglycerol headgroup; OL: oleate acyl chain; PA: palmitate acyl chain. **(C)** Lateral pressure profile (Π) of the simulated bilayers, showing the characteristic negative component associated with the membrane:solvent interface (see the water density drop in B) and a central positive pressure component in the POPC:POPG system (top) that is absent in the DOPC:DOPG bilayer [23]. PL and PN are lateral and normal components of the pressure tensor. **(D)** Distributions of the area-per-lipid in the simulated bilayers. The area per lipid is significantly higher (p < 0.001; Tukey HSD test, Table 1) in the bilayer composed of DOPC:DOPG, despite overall similar electron density profiles. **(E)** 2D number density of lipids on the x z plane of the POPC:POPG bilayer and the difference of the x z densities and lateral pressure profiles of (POPC:POPG – DOPC:DOPG). In the DOPC:DOPG bilayer, the density is higher in the membrane center and in the interfacial regions but lower in the region of the unsaturation and the headgroups. In the pressure profile, two UFAs cause a higher repulsive component in the head group region (1), a more attractive pressure close to the water membrane interface (2), a higher repulsive component in the region corresponding to the double bonds (3), and a lower repulsion in the center of the membrane bilayer (4). The regions with a more attractive component in the DOPC:DOPG system (2,4) relate to a relatively higher density of lipids. Δ9 indicates the region where the unsaturated bonds in the upper leaflet of the lipid bilayer are located (see B, green curves). All values from B to E were calculated from all-atom MD simulations based on five independent replicas.

To scrutinize the lipid organization within the membrane at the molecular level, we carried out all-atom molecular dynamics (MD) simulations of DOPC:DOPG and POPC:POPG lipid bilayers. Both systems showed very similar electron density profiles and membrane thicknesses (Figure 3B and Suppl. Figures 3; Table 1). However, a comparison of the lateral pressure profiles and the lipid packing revealed prominent differences between these lipid bilayers (Figure 3C-E). Pressure differences were observed in 1) the head group region (15-17 Å, the repulsive component is stronger in DOPC:DOPG); 2) close to the ester bonding (11-14 Å, attractive pressure in DOPC:DOPG); 3) in the region of the unsaturation (~5-10 Å, the repulsive component is stronger in DOPC:DOPG); and 4) in the membrane center (0 Å, only the POPC:POPG system has a repulsive component). Thus, the presence of an additional double bond in DOPC:DOPG membrane shifts repulsive pressure from the acyl chain region towards the water interface, as suggested by analytical studies and found for similar lipid compositions of varying unsaturation degrees [22,23], and to some extent to the head group region. For DOPC:DOPG, the more attractive pressure in the interface and at the membrane center comes at the expense of the stronger repulsive components in the regions of the head groups and the unsaturation, reflecting more pronounced steric interactions between the polar groups and the acyl chains, respectively. Indeed, the packing density in the DOPC:DOPG system is higher in the interface region and the membrane center, but lower in the regions of the head groups and the unsaturation (Figure 3E), similar to what has been previously found for pure DOPC versus POPC systems [13]. The more heterogeneous local number density is associated with a larger area per lipid of the DOPC:DOPG system (70.25 ± 0.05 Å^2^, mean ± SEM) compared to the POPC:POPG system (66.74 ± 0.02 Å^2^; Figure 3D and Table 1), in agreement with the results from laurdan fluorescence (Figure 3A). Thus, both the experimental test and MD simulations indicate that the elevated UFA content leads to redistribution of the pressure within the membrane and induces looser packing of the lipid head groups.

**Table 1.** Area per lipid and membrane thickness measured in MD simulations of the investigated systems.^a^

| Head group | POPC:POPG<br>70:30 | DOPC:DOPG<br>70:30 | POPC:POPG:<br>bis(PO)CL<br>70:25:5 | POPC:POPG:TOCL<br>70:25:5 | POPC:bis(PO)CL<br>85:15 | POPC:TOCL<br>85:15 |
| --- | --- | --- | --- | --- | --- | --- |
| **PC APL<sup>b,†</sup>** | 67.14 (0.05) | 70.68 (0.07) | 68.94 (0.18) | 69.12 (0.16) | 73.05 (0.12) | 74.13 (0.12) |
| **PG APL<sup>b,†</sup>** | 65.86 (0.19) | 69.28 (0.18) | 67.12 (0.09) | 67.49 (0.20) | - | - |
| **CL APL<sup>b,†</sup>** | - | - | 77.72 (1.30) | 78.30 (1.04) | 79.23 (0.65) | 80.38 (0.23) |
| **Average APL<sup>c,†</sup>** | 66.74 (0.02) | 70.25 (0.05) | 68.88 (0.08) | 69.13 (0.10) | 73.97 (0.09) | 75.06 (0.07) |
| **Thickness<sup>d</sup>** | 37.38 (0.01) | 37.06 (0.02) | 37.86 (0.04) | 37.90 (0.05) | 38.71 (0.04) | 38.64 (0.04) |

<sup>a</sup> All measurements were done over the last 800 ns of five replicas and are shown as the mean of replica means (standard error of the mean).<sup>b</sup> Area per lipid ( $\text{\AA}^2$ ) as measured by APL@Voro [66].<sup>c</sup> Area per lipid ( $\text{\AA}^2$ ) measured as the xy-sectional area of the average simulation box used per lipid.<sup>d</sup> Membrane thickness ( $\text{\AA}$ ) measured as the distance between the z-coordinates of the centers of mass of the phosphorous atoms of each leaflet.<sup>†</sup>The pairwise difference of the mean across all systems is statistically significant ( $p < 0.001$ , Tukey HSD).

### Unsaturated fatty acids facilitate SecA binding to the lipid bilayer

SecA:lipid interactions at the membrane interface are recognized as an essential factor in Sec-mediated translocation [3,5,24]. Peripheral SecA:lipid binding may be a prerequisite to activate the ATPase and ensure the downstream SecA:SecYEG assembly [7,25]. The N-terminal tail of SecA (residues 1-25) forms an amphipathic helix that binds to and likely sinks into the lipid membrane in the presence of anionic head groups due to electrostatic interactions (Figure 4A). In unbiased MD simulations of 1 μs length, the SecA N-terminal tail interacts with DOPC:DOPG and POPC:POPG lipid head groups mainly through its basic residues (Lys-4, Lys-8, Arg-13, Arg-16, Arg-19, Arg-20 and Arg-22; Figure 4B and Suppl. Figure 4B). Deleting the N-terminal tail abolishes SecA binding to liposomes and inhibits the preprotein translocation (Suppl. Figure 4A). However, the activity can be restored *in vitro* once the histidine-tagged SecA is artificially anchored to proteoliposomes [26], so the primary role of the N-terminus is to facilitate SecA binding to the membrane.

**Figure 4.**
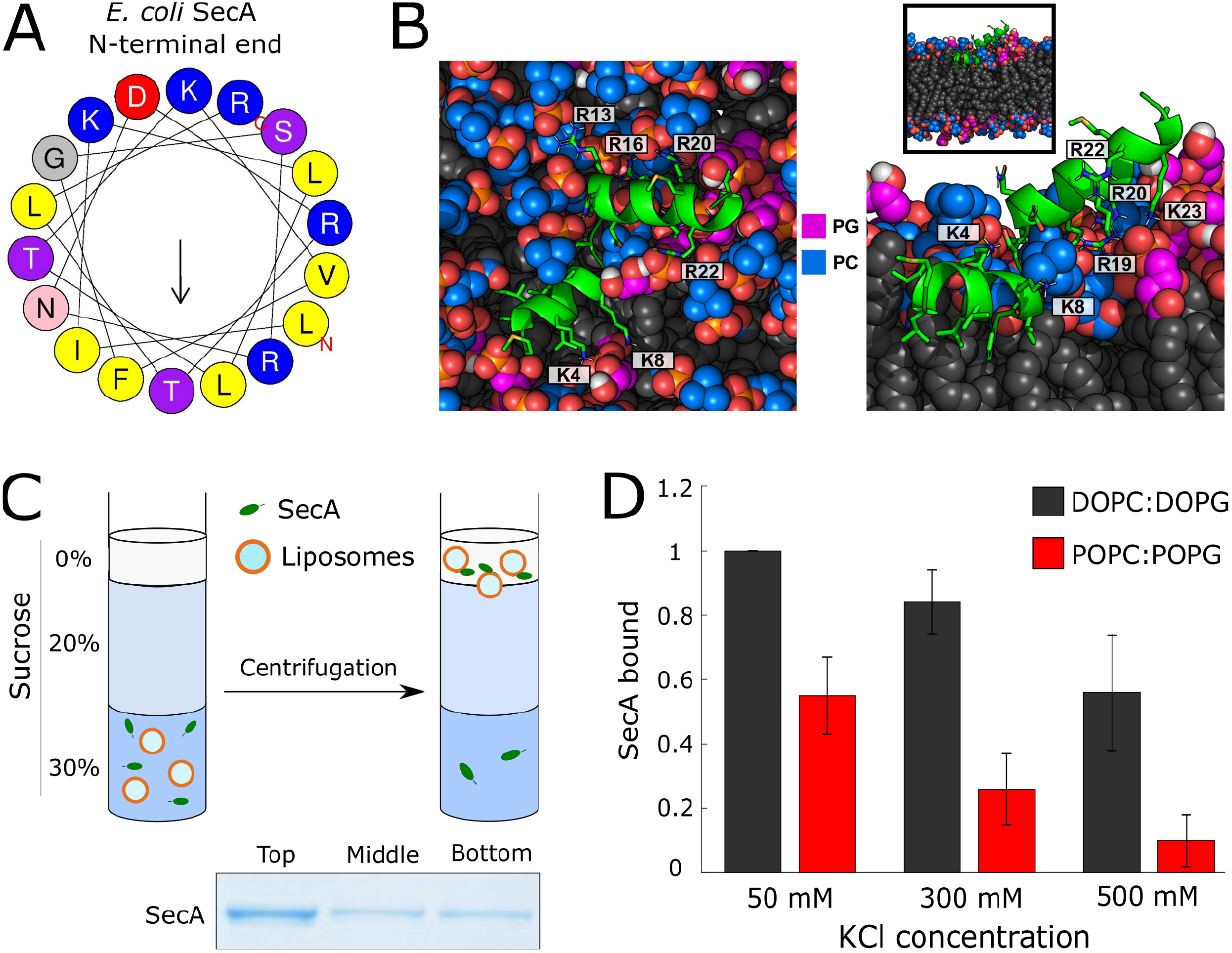
SecA:membrane association is mediated by unsaturated fatty acids. **(A)** Helix wheel projection of the N-terminal end of *E. coli* SecA of the putative amphipathic helix (via heliquest.ipmc.cnrs.fr). The hydrophobic moment (~0.4) is indicated with the arrow. **(B)** A binding pose of the N-terminal amphipathic helix of SecA at the interface of the lipid bilayer after 1 μs of MD simulation (DOPC:DOPG, replica 4). Basic residues that show pronounced interactions with the head groups are highlighted (Suppl. Figure 4B). **(C)** Scheme of the flotation assay to probe SecA:lipid interactions. Upon the centrifugation, the liposomes loaded with aqueous buffer migrate to the top of the sucrose density gradient. Liposome-bound SecA is found in the top layer of the gradient, while free SecA remains in the bottom fraction. **(D)** The flotation assay reveals UFA-dependent and salt-sensitive binding of SecA to the liposomes.

Although the effect of UFAs on SecA binding has not yet been examined, it has been shown for several eukaryotic proteins that defects in the lipid packing and transiently exposed hydrophobic areas enhance the affinity of amphipathic helices to membranes [12,14]. To test whether the UFA content and the altered lipid packing affect the SecA:membrane interaction, binding of SecA to liposomes was examined. Once bound to the lipid leaflet, SecA can float with liposomes through a sucrose density gradient, thus allowing to estimate the binding efficiency (Figure 4C). SecA readily interacted with PG-containing liposomes, and the binding was salt-sensitive, as expected for the electrostatics-driven interaction [3] (Figure 4D). Increasing the salt concentration from 50 to 500 mM reduced SecA binding to POPC:POPG liposomes by ~80 %. Notably, for DOPC:DOPG liposomes the reduction was limited to 40 % only. Over the whole range of tested salt concentrations, the amount of SecA bound to DOPC:DOPG liposomes exceeded that bound to POPC:POPG. The difference reached 7-fold at 500 mM KCl, thus confirming the effect of the lipid packing on SecA:membrane interaction. Binding of SecA to gel-phase membranes DPPC:DPPG was suppressed three-to four-fold even at the low salt concentration (50 mM KCl; Suppl. Figure 4C), despite the presence of the anionic lipids.

Notably, when POPC was substituted with DOPC in a stepwise manner, both SecA binding and the preprotein translocation in SecYEG-proteoliposomes continuously increased, and introducing DOPG instead of POPG had a similar effect (Suppl. Figure 5A and 5B). This clear correlation supported the hypothesis that the UFA-enhanced SecA:lipid interactions promote SecA:SecYEG translocation. We questioned then whether the position or configuration of the double bond within the UFA affects SecA:membrane interactions. Placing *cis*-bonds in position Δ6 of PC lipids rendered a loosely packed membrane interface, as it was confirmed by laurdan fluorescence polarization (GP = −0.510±0.004), and the liposomes manifested highly efficient SecA binding (Suppl. Figures 5C) and stimulated translocation in presence of SecYEG (Suppl. Figure 5D). *Trans*-bonds in both fatty acids of PC had an opposite effect on the membrane properties: Although the lipid bilayer resided in the fluid phase (phase transition temperature ~12 °C), the laurdan fluorescence indicated tight packing within the head group region similar to UFA-poor POPC:POPG membranes (Figure 3A). The increase in the lipid packing led to suppressed binding of SecA and finally caused 10-fold reduction in SecA:SecYEG-mediated translocation (Suppl. Figure 5C and 5D).

**Figure 5.**
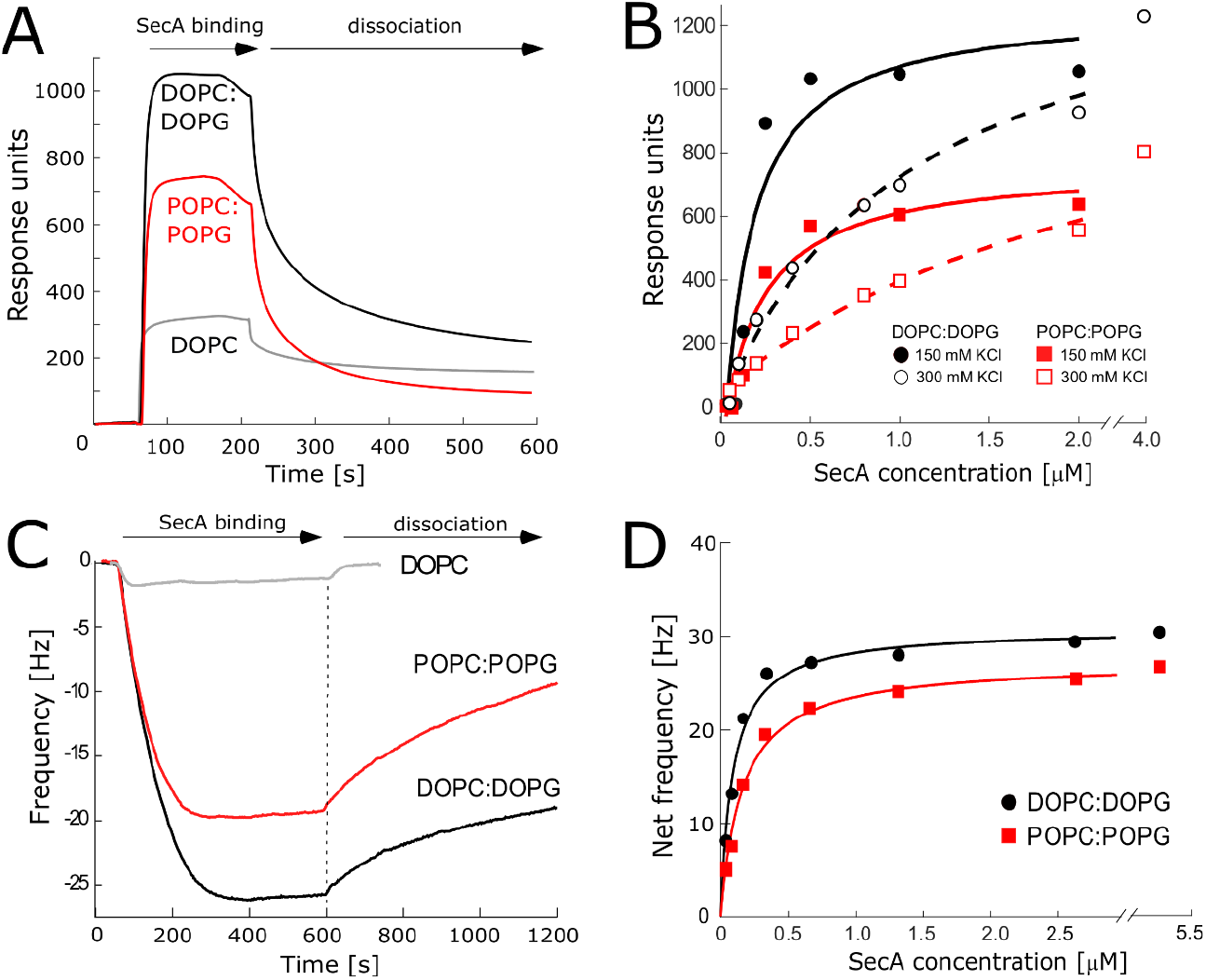
Unsaturated fatty acids enhance the affinity of SecA:membrane interactions. **(A)** Surface plasmon resonance (SPR) sensograms of SecA binding to liposomes, followed by the dissociation phase. Charge-neutral DOPC membranes were used as a negative control. SecA concentration: 500 nM. **(B)** Analysis of the steady-state SPR response over a range of SecA concentrations (31 nM to 2 μM) reveals enhanced binding to DOPC:DOPG membranes at varying salt concentrations. **(C)** Quartz crystal microbalance sensograms of SecA binding to planar lipid bilayers, followed by the dissociation phase. Charge-neutral DOPC membranes were used as a negative control. SecA concentration: 800 nM. (D) Analysis of the steady-state frequency change over a range of SecA concentrations (40 nM to 5.25 μM) reveals enhanced binding of SecA to DOPC:DOPG membranes

Aiming for quantitative characterization of SecA:lipid interactions, surface plasmon resonance (SPR) experiments were carried out. The steady-state response upon binding of SecA to the chip-anchored liposomes was strongly enhanced in UFA-based DOPC:DOPG membranes (Figure 5A). The measurements over a range of SecA concentrations provided an estimate of the apparent dissociation constant *K*_D_ for SecA:lipid interactions (Figure 5B). In the presence of 150 mM KCl and 5 mM MgCl_2_, SecA showed a 1.7-fold higher affinity to DOPC:DOPG membranes (125±5 nM) than to POPC:POPG membranes (210±45 nM). In agreement with the flotation assay (Figure 4D), SecA:lipid binding was suppressed at elevated salt concentration of 300 mM KCl for both DOPC:DOOG and POPC:POPG, where the apparent *K*_D_ decreased to ≃ 1.2 μM and 2.4 μM, respectively (Figure 5B).

To exclude that binding was affected by the non-physiological positive curvature of the liposomes, a complementary experiment was carried out using planar supported lipid bilayers (SLBs) deposited on a quartz crystal microbalance (QCM) chip [27]. Liposomes were non-covalently adsorbed on the quartz chip surface and fused to form continuous SLBs (Suppl. Figure 6). Subsequent SecA binding increased the mass adsorbed on the quartz chip, which affected its resonance frequency (Figure 5C). Depending on SecA concentration, the ATPase binding to DOPC:DOPG membranes measured in 150 mM KCl and 5 mM MgCl_2_ was 25 to 50% higher than to POPC:POPG membranes, and the apparent *K*_D_ values were 91 nM and 161 nM, respectively (Figure 5D), in a good agreement with the SPR data. Thus, biochemical and biophysical assays confirmed the differential binding of SecA to lipid bilayers depending on the UFA content and the associated lipid head group packing, and the binding efficiency correlated with the translocation activity of the SecA:SecYEG complex.

### Tetraoleoyl-cardiolipin stimulates SecA binding and preprotein transport

The inner membranes of *E. coli* and other Gram-negative bacteria commonly contain a minor fraction of cardiolipin (CL) molecules [15], and a recent mass-spectrometry analysis revealed that the most abundant CL species in the *E. coli* membrane contain four mono-UFAs [19]. CL molecules are enriched two-to three-fold in the proximity of the translocon [8,19], and it has been suggested that CL facilitates SecYEG homo-dimerization, but may also serve as an acceptor for protons to contribute to the proton-motive force [8]. However, the functional translocon *in vitro* and *in vivo* is built of monomeric SecYEG, and no stimulatory effect of CL on SecA:SecYEG activity in the UFA-enriched membranes was observed [9,28]. Thus, the role of CL in SecA:SecYEG activity has remained intensively disputed [7,29].

In light of the discovered effect of UFAs on Sec-mediated translocation, we questioned whether UFA-enriched CL, such as tetraoleoyl-CL (TOCL), may recruit SecA to the lipid membrane and to enhance the preprotein transport. To test this hypothesis, SecYEG was reconstituted in POPC:POPG:TOCL membranes with a variable amount of TOCL. To keep the net negative charge at the membrane interface constant, the variations in the CL content were compensated by tuning the POPG fraction. Changing the CL fraction from 0 to 15 mol % increased the translocation activity up to 10-fold, indicating that TOCL is indeed a potent stimulator of protein translocation (Figure 6A). To exclude potential dimerization of SecYEG in the presence of CL, the ATTO 643-labeled translocons were reconstituted into nanodiscs as monomers, and the translocation efficiency in the presence of TOCL was determined using the FRET-based assay [30,31]. The dimensions of nanodiscs (outer diameter ~15 nm) and the lipid:protein ratio used upon the reconstitution ensured that ~200 lipid molecules were embedded in each nanodisc [32]. TOCL strongly stimulated the translocation efficiency and kinetics, despite the nanodisc boundaries physically prevented dimerization of SecYEG (Figure 6B). While TOCL did not affect the stability of SecYEG (*T*_m_ ~ 66 °C, Figure 2C), binding of SecA to TOCL-containing liposomes was strongly enhanced compared to POPC:POPG, as tested by flotation assay and SPR (Figure 6C and Suppl. Figure 7A). Although additional effects of interactions of TOCL with the active SecA:SecYEG complex are possible, our data demonstrate that even in the absence of the proton motive force TOCL favours SecA:lipid interactions at the membrane interface and stimulates the translocation.

**Figure 6.**
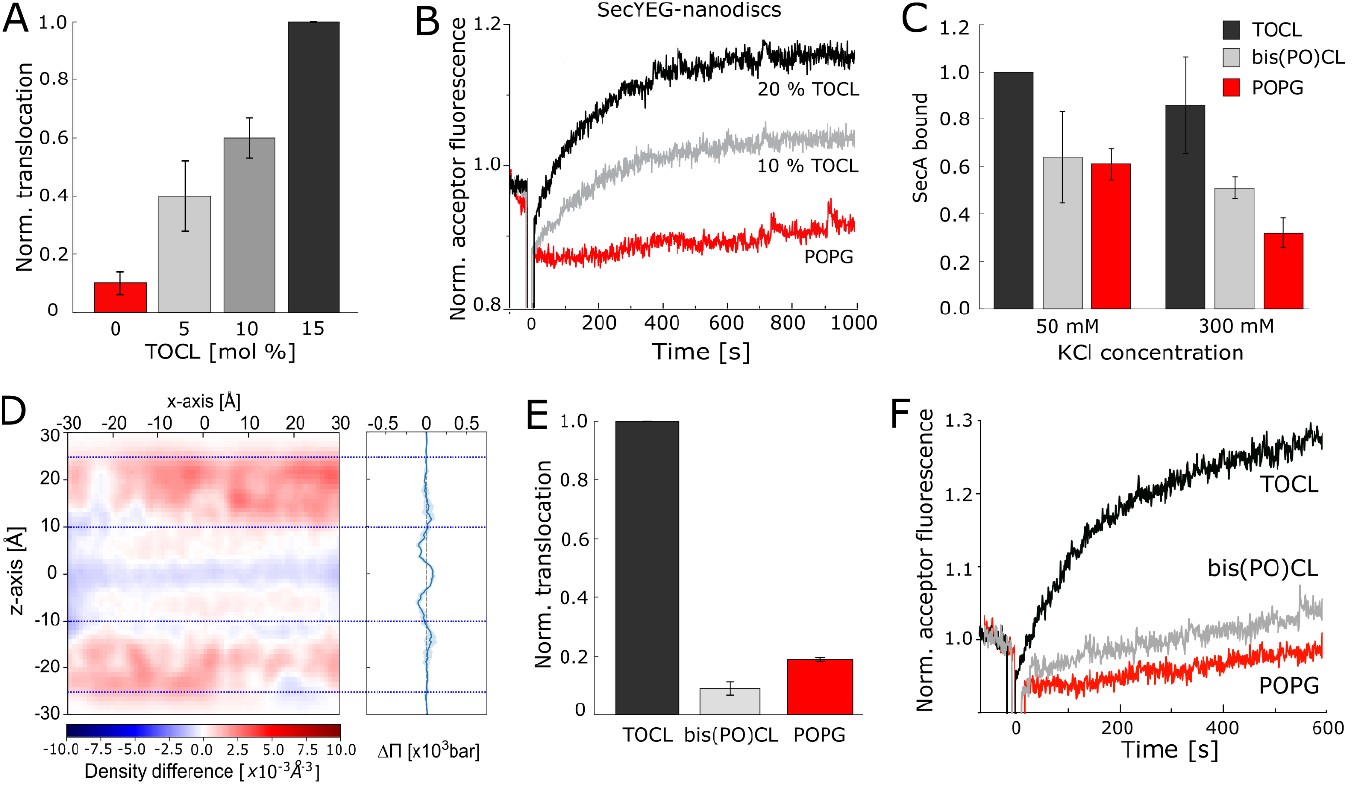
UFA-enriched cardiolipin stimulates Sec-mediated translocation. **(A)** Elevated concentrations of tetraoleoyl-cardiolipin (TOCL) stimulate preprotein transport into liposomes. **(B)** FRET-based translocation assay shows increased activity of monomeric SecYEG in nanodiscs in presence of TOCL. The stimulated activity is achieved without oligomerization of the translocon. **(C)** Flotation assay confirms the preferential interactions of SecA with UFA-enriched TOCL. Binding to TOCL-containing membranes is the least affected by the elevated ionic strength, so the interactions with UFAs promote the membrane-bound form of SecA. **(D)** The difference in 2D density profile and lateral pressure obtained by MD simulations for POPC:CL lipid bilayers (bis(PO)CL-TOCL) reveal a qualitatively similar trend as that observed for the POPC:POPG – DOPC:DOPG case (Figure 3E). UFA-specific SecA:membrane interactions in the presence of TOCL correlate with the translocation efficiency (E) and the translocation rate probed in FRET-based assay (F).

### Effect of cardiolipin on translocation depends on the fatty acid structure

Does the CL-mediated stimulation originate from the UFA content, or may it be determined by the high charge density within the CL head group? To address this question, two CL variants, TOCL (four UFAs) and bis(palmitoyloleoyl)-CL (bis(PO)CL, two UFAs), were examined with MD simulations and in model liposomes. Exchanging 5% of POPG with either TOCL or bis(PO)CL in the simulated POPC:POPG bilayer caused significant increases in the overall and lipid type-specific area per lipid with respect to the POPC:POPG system (Table 1; Suppl. Figures 3 and 8). To test whether increasing the cardiolipin fraction causes more pronounced effects, we also evaluated systems with POPC:bis(PO)CL and POPC:TOCL (molar ratio 85:15). In both cases a significant increase in the average area per lipid was observed, and the change was the most pronounced for the POPC:TOCL bilayer (Table 1). Given the larger size of cardiolipins, an increase in the average area per lipid was to be expected, but, notably, the effect also comprised the other lipid types in the bilayer, so the area per POPC molecule increased by 10 %, from 67 Å^2^ (POPC:POPG bilayer) to 74 Å^2^ (POPC:TOCL bilayer). Next, we analysed the distribution of the number density of the simulated bilayers with TOCL or bis(PO)CL. The density difference suggested that the TOCL causes a higher density in the interface region and lower density in the head group region than bis(PO)CL (Figure 6D), being in a qualitative agreement with the results acquired from the simulated DOPC:DOPG and POPC:POPG bilayers (Figure 3). Thus, both CL types caused substantial re-structuring of the lipid bilayer compared to the POPC:POPG system, as indicated by the change in the lipid packing and the transmembrane distribution of the pressure density.

Notably, the density difference calculated between the bilayers with TOCL and bis(PO)CL is smaller in magnitude than the value determined for pure DOPC:DOPG and POPC:POPG membranes. In agreement with the computed predictions, the laurdan fluorescence in CL-containing liposomes manifested a modest, although significant, change in the polarization value upon variations in the UFA content: The general polarization decreased from 0.389 ± 0.005 (15 mol % TOCL) to 0.335 ± 0.012 (15 mol % bis(PO)CL). To probe the effect of different CLs on SecA:SecYEG activity, functional tests were carried out using POPC:CL liposomes. SecA:lipid interactions were enhanced by TOCL, but not bis(PO)CL species in comparison to pure POPC:POPG membranes (Figure 6C), suggesting that the UFA abundance, but not the charge density within the CL, affect the ATPase binding. Furthermore, the UFA-dependent binding of SecA converted into the differential effect of CLs on preprotein transport: While no stimulation was provided by bis(PO)CL, 15 mol % TOCL stimulated the activity of SecA:SecYEG (Figure 6D), and the translocation efficiency correlated with the ATP hydrolysis rates (Figure 6E). The FRET-based translocation assay further confirmed a prominent effect of TOCL, but not bis(PO)CL on the translocon activity (Figure 6F), so the composition of the membrane core influenced the transport rate of a polypeptide chain.

## Discussion

The diversity of lipids found in prokaryotic and eukaryotic cellular membranes greatly determines physicochemical properties of the bulk membrane, but also promotes formation of functional membrane domains and mediates specific interactions with the membrane-associated proteins [33,34]. Both bulk and specific local properties of the lipid bilayer may regulate the functionality of membrane-embedded complexes, being crucial for cellular pathways, such as energy metabolism, signal transduction, and the broad repertoire of transport processes. Protein translocation via the essential Sec machinery is a well-known example, where the naturally abundant anionic lipids are required for the assembly of the functional SecA:SecYEG complex [3,7,25]. While the contribution of electrostatic interactions has been extensively examined, here we reveal that the protein translocation is sensitive to changes in the fatty acid composition of the lipid bilayer. The protein translocation is augmented manifold upon increasing the content of dioleoyl fatty acids in the range from 50 to 100 mol %, although the fluidity of the bilayer is not changed. While neither stability nor topology of the translocon SecYEG are affected by the lipid composition, the elevated translocation correlates with the increased binding of the ATPase SecA to the UFA-enriched lipid bilayer. The stimulatory effect of UFAs on SecA binding is well-pronounced at the elevated salt concentrations, pointing to the previously not appreciated role of hydrophobic interactions.

SecA:lipid binding is largely mediated by the N-terminal amphipathic helix of the ATPase: The helix is essential for interactions with the membrane, where it extensively binds anionic lipids via lysine and arginine residues at the polar side [26,35,36]. Further partitioning of the helix into the lipid leaflet, however, must rely on interactions with the hydrophobic fatty acids. Our all-atom MD simulations together with biophysical analysis of the lipid bilayer structure suggest that UFAs induce irregularities and looser head group packing at the membrane:solvent interface. A similar effect must have been observed in the early study on SecA:SecYEG, when DOPE was described as a potent stimulator of the translocon machinery [5]. The small PE head group combined with two UFA chains results in extensive packing defects at the membrane interface [12]. In the simplest scenario, those interfacial features allow the amphipathic helix of SecA to access the transiently exposed hydrophobic regions and facilitate partitioning of the helix into the bilayer [12,24,36]. At the elevated salt concentrations, the electrostatic coupling of the protein to the anionic lipids largely deteriorates, and the membrane-bound state of SecA depends on non-polar contacts with the membrane interior.

Once bound to the lipid interface, SecA undergoes two-dimensional diffusion within the lipid leaflet prior docking on the SecYEG translocon [25,37]. Thus, the elevated concentration of the membrane-bound SecA favours the downstream assembly of the SecA:SecYEG complex and results in the immense stimulation of preprotein transport, as observed in our experiments. Notably, the quantitative analysis of SecA:lipid interactions via SPR and QCM reveals a modest 2-to 3-fold increase in the affinity once the UFA content reaches 100 mol %, while the translocation activity of the SecA:SecYEG complex is stimulated up to 7-fold. This may indicate that the preprotein translocation involves multiple SecA molecules, which diffuse at the membrane interface and transiently bind the translocon, either as dimers or monomers, to perform cycles of translocation. Despite the extensive research, neither the quaternary state and dynamics of SecA nor its processivity in translocation have been clarified [38–40]. Recent reports have suggested that the oligomeric state of the membrane-bound SecA depends on the membrane lipid composition [35,41], so it deserves further evaluation, whether UFA-mediated lipid packing influences SecA dynamics or, potentially, diffusion at the membrane interface.

The requirement of CL for SecA:SecYEG-mediated translocation has been intensively debated, and specific SecYEG:CL and SecA:CL contacts and CL-induced protein dimerization, which promote the preprotein transport, have been proposed [29,42,43]. Lately, the stimulatory effect of CL has been described in presence of the proton motive force [8]. Here, we demonstrate that the UFA-enriched TOCL, but not bis(PO)CL, contributes to SecA binding and enhances the translocation up to 10-fold. Furthermore, the translocation experiments in nanodiscs allowed to rule out the translocon dimerization upon the translocation. Thus, UFA-mediated SecA binding does not depend on lipid species, but strongly correlates with the bulk head group packing at the membrane interface. Interestingly though, MD simulations and measurements of the laurdan fluorescence reveal relatively small differences between lipid membranes containing TOCL or bis(PO)CL, while only TOCL is potent to stimulate the translocation. It suggests that more intricate mechanisms should be considered to interpret the effect of UFAs on SecA binding and translocation, e.g., formation of CL-specific domains within the otherwise miscible lipid membrane [44], restructuring of the annular lipids upon SecA binding, or UFA-dependent dynamics of the SecA:SecYEG complex. Further experiments, both in model systems and native bacterial membranes, and extensive MD simulations of protein:lipid interfaces will aim to explain the effect of the membrane structure on SecA:SecYEG activity.

The demonstrated role of UFAs as mediators of SecA binding and protein translocation may be a critical factor for the reaction in living cells. Bacteria tune their membrane lipid composition in response to environmental factors and growth conditions, so the fraction of mono-UFAs in mesophilic bacteria, such as *E. coli* and *Pseudomonas aeruginosa*, may vary from 50 to 75 mol % [10,11,45]. Low temperatures are associated with the elevation of the UFA content, either via *de novo* lipid synthesis or the enzymatic desaturation of existing fatty acids within the membrane. Our data suggest that those physiological modifications will favour the membrane-bound state of SecA and thus promote protein transport to compensate for the temperature-dependent kinetics decay. Complementary, under conditions of hyperosmotic shock and increased intracellular salt concentration or dissipation of the proton motive force, the hydrophobic interactions of SecA with unsaturated lipids will prevent dissociation of the ATPase from the membrane. Likewise, it is to expect that SecA homologs from extremophile species are evolutionary tuned for interactions with UFA-depleted membranes, e.g. via the strong hydrophobic dipole at the N-terminal domain. Comparative functional analysis of SecA homologs from those species, and potentially the reconstituted SecA:SecYEG machinery, may further advance the understanding of the molecular adaptation mechanisms.

## Materials & methods

### Protein purification

Overexpression of SecYEG with the N-terminal deca-histidine tag in *E.coli* C41(DE3) cells was induced with 0.5 mM IPTG and carried out for 2 hours at 37 °C. Cells were harvested by centrifugation and resuspended in buffer R (50 mM KOAc, 20 mM Hepes/KOH pH 7.4, 5 mM Mg(OAc)_2_, 5 % glycerol, 1 mM DTT and cOmplete Protease inhibitor cocktail (Roche). Cells were lysed (Microfluidizer, M-110P, Microfluidics Corp.) and the debris was removed by centrifugation. Crude membranes were pelleted by centrifugation for 1 h at 40,000 rpm (45 Ti rotor, Beckman Coulter) and then resuspended in buffer R. The membranes were solubilized using 1 % DDM in presence of 500 mM KCl, 50 mM Hepes/KOH pH 7.4, 5 % glycerol, 200 μM TCEP, and the protease inhibitor cocktail. His_10_-tagged SecYEG was isolated using Ni^2+^-NTA agarose resin (Qiagen). Once bound to Ni^2+^-NTA beads, the single-cysteine SecY^C148^EG variant was optionally labelled with ATTO 643-maleimide (ATTO-Tec GmbH) upon incubation with 100-200 μM dye for 2 h at 4 °C [9]. SecYEG-loaded resin was extensively washed with 50 mM Hepes/KOH pH 7.4, 500 mM KCl, 5 % glycerol, 10 mM imidazole, 200 *μ*M TCEP, 0.1 % DDM, and the protein was eluted with the buffer containing 300 mM imidazole. The protein was transferred to 50 mM Hepes/KOH pH 7.4, 150 mM KCl, 5 % glycerol, 0.05 % DDM, 200 *μ*M TCEP using PD MidiTrap G-25 column (Cytiva/GE Life Sciences). Optionally, size exclusion chromatography (SEC) was carried out using Superdex 200 30/10 GL Increase column (Cytiva/GE Life Sciences) to control the homogeneity of the sample (elution peak at 12.5 mL corresponds to SecYEG in a DDM micelle, total mass of approx. 145 kDa). Protein concentration was determined spectrophotometrically (extinction coefficient 72,000 M^-1^ cm^-1^). The expression and purification steps were controlled with SDS-PAGE (Quick Coomassie stain, Serva), and in-gel fluorescence of the SecY-conjugated ATTO 643 dye was visualized (Amersham Imager 680RGB, GE Healthcare Life Sciences).

SecA gene was cloned into pET21a plasmid to carry the C-terminal His6-tag. SecA overexpression in *E. coli* BL21(DE3) cells was induced with 0.5 mM IPTG and carried out for 2 hours at 30 °C. Cells were then harvested by centrifugation and resuspended in 50 mM KOAc, 20 mM Tris/HCL pH 7.5, 5 mM Mg(OAc)_2_, 20 % glycerol, 1 mM DTT, and cOmplete protease inhibitor cocktail. Cells were lysed and the lysate was clarified by centrifugation for 1 h at 100,000g (45 Ti rotor, Beckman Coulter). The clarified lysate was incubated with Ni^2+^-NTA agarose resin for 1 h at 4 °C, the resin was then washed with 500 mM KOAc, 20 mM Tris/HCL pH 7.5, 5 mM Mg(OAc)_2_, 20 % glycerol, 20 mM imidazole, 1 mM DTT. SecA was eluted with the buffer containing 300 mM imidazole. The eluted fractions were concentrated using Amicon Ultra-4 filtering device, cut-off size 50 kDa (Millipore), and SecA was subject to SEC using Superose 6 Increase 10/300 GL column (Cytiva/GE Life Sciences) in 50 mM KOAc, 20 mM Tris/HCL pH 7.5, 5 mM Mg(OAc)_2_, 20 % glycerol, 1 mM DTT, resulting in a peak at ~15.5 mL elution volume. Peak fractions were pooled together and the protein concentration was determined spectrophotometrically (extinction coefficient 75,000 M^-1^ cm^-1^). Precursor proteins proOmpA and proOmpA-DHFR were overexpressed in inclusion bodies as previously described elsewhere [46,47] and stored in 8 M urea.

### Liposomes preparation

Synthetic lipids and *E. coli* polar lipid extract were purchased from Avanti Polar Lipids, Inc. as stocks in chloroform. The lipids were mixed in desired ratios, and chloroform was evaporated under vacuum conditions at 40 °C using a rotary evaporator (IKA). The dried lipid film was then rehydrated in 50 mM KCl, 50 mM Hepes/KOH pH 7.4 to achieve the lipid concentration of 5 mM. To form large unilamellar vesicles the crude liposomes were manually extruded through the porous polycarbonate membranes (Nuclepore, Whatman) using the Mini-Extruder set (Avanti Polar Lipids, Inc.). The membrane pore size was decreased stepwise from 400 nm to 200 nm and vesicle sizes were controlled by dynamic light scattering (Nicomp 3000, Entegris, Inc.).

### SecYEG reconstitution

SecYEG variants were reconstituted in liposomes with different composition at 1:1,000 protein:lipid ratio as follows. Liposomes were swelled by adding 0.2 % DDM followed by incubation for 10 minutes at 40 °C, and then mixed with SecYEG in 0.05 % DDM. The reconstitution mixture was incubated for 30 minutes on ice. The detergent was then removed upon incubation with Bio-Beads SM-2 sorbent (Bio-Rad Laboratories) overnight at 4 °C. Formed proteoliposomes were pelleted upon centrifugation at 80,000 rpm for 30 minutes (S120-AT3 rotor, Thermo Fisher/Sorvall) and resuspended in 50 mM KCl, 50 mM Hepes/KOH pH 7.4 to achieve the final translocon concentration of 5 *μ*M. To examine the reconstitution efficiency, the proteoliposomes were subjected to centrifugation in the density gradient. 100 μL of proteoliposomes were diluted and mixed with 54 % iodixanol-based medium (Optiprep™, Merck/Sigma) to achieve the final iodixanol concentration of 40 %. 30 % iodixanol was then layered on top followed by 15 % and 5 %. Samples were centrifuged for 3 h at 200,000 g (TST 60.4 swing-out rotor, Thermo Fisher/Sorvall). The gradients were manually fractionated, the contents were precipitated with trichloroacetic acid (TCA) and analysed on SDS-PAGE.

SecYEG topology in the formed proteoliposomes was analysed by probing the accessibility of the SecY cytoplasmic side. The recombinant SecYEG bears the N-terminal histidine tag on SecY subunit followed by an enterokinase cleavage site [5]. By cleaving the tag from SecYEG reconstituted in liposomes, the amount of correctly oriented SecYEG could be determined due to difference in migration on SDS-PAGE. 5 μL proteoliposomes were mixed with 8 units of enterokinase light chain (New England Biolabs) and diluted to 20 μL, then samples were incubated at 25 °C overnight.

Reconstitution of SecYEG into MSP2N2 nanodiscs was carried out following the previously established protocol [6,30,32]. To achieve the monomeric state of the translocon, a large excess of lipids and the scaffold protein MSP was supplied into the reconstitution reaction (SecYEG:MSP:lipid ratio of 1:10:1,000). SecYEG-loaded and empty nanodiscs were separated via size exclusion chromatography using Superdex 200 10/300 Increase GL column (Cytiva/GE Life Sciences).

### *In vitro* translocation assay

The previously described protocol for the fluorescently labelled preprotein translocation *in vitro* was followed with minor modifications [9,48]. 10 μL of proteoliposomes was mixed with 1 μM SecA (concentration for monomer), 0.5 μM SecB (concentration for tetramer), energy mix (0.05 mg/mL phosphocreatine kinase, 10 mM phosphocreatine), 0.1 mg/mL BSA, 10 mM DTT, 10 mM MgCl_2_, and 1 *μ*M proOmpA labelled with fluorescein-maleimide (proOmpA-FM). The total volume was adjusted to 50 *μ*L with buffer (50 mM KCl, 50 mM Hepes/KOH pH 7.4) and samples were equilibrated at 37 °C for 5 min. The translocation was triggered by the addition of 6 mM ATP and left to proceed for 15 minutes. Afterwards, 10 % volume was withdrawn as a reference for the input of proOmpA-FM, and 0.2 mg/mL proteinase K was added to the remaining sample to degrade the non-translocated preprotein. After incubation for 15 min, 150 *μ*L 20 % TCA was added, and the aggregated samples were pelleted via centrifugation at 20.000 g for 10 min (Eppendorf 5417 R table-top centrifuge). The pellets were washed with 500 *μ*L ice-cold acetone and centrifuged for 5 min. Acetone was discarded and the dried pellets were resuspended in 15 *μ*L 2.5x SDS-PAGE sample buffer, incubated for 5 minutes at 95 °C and then loaded on SDS-PAGE. In-gel fluorescence of the protease-protected proOmpA-FM was recorded and quantified (ImageQuant TL, Cytiva/GE Life Sciences). The background signal was subtracted using the implemented *Local average* algorithm. The amount of the transported proOmpA-FM was determined based on the available reference sample. At least two independent tests were carried out for each experiment.

### FRET-based real time kinetics assay

FRET-based analysis of the preprotein translocation was carried out following the previously established protocol [9,25]. Briefly, proOmpA^C292^-DHFR fusion protein was labelled with Cyanine3-maleimide (donor) at the unique cysteine at position 292 of proOmpA domain, and SecY^C148^EG was labelled with ATTO 643-maleimide (acceptor) at the periplasmic side. To record the acceptor fluorescence, the monochromators of the thermostated spectrofluorometer (Fluorolog-3, Horiba™ Scientific) were set to the excitation wavelength of 510 nm (slit width 3 nm) and the emission wavelength of 690 nm (3 nm slit width). Prior the translocation, the DHFR domain was stably folded in presence of the co-factors methotrexate and NADPH, as previously described [9]. 100 nM proOmpA-DHFR was mixed with 5 μL proteoliposomes (400 nM SecYEG, dual orientation) in presence of 1 μM SecA, and completed to 60 μL of buffer (150 mM KCl, 5 mM MgCl_2_, 50 mM Hepes/KOH pH 7.4). The quartz cuvette (Hellma Analytics) was incubated at 37 °C for 150 sec and the translocation reaction was triggered by the addition of 5 mM ATP. The acceptor fluorescence was recorded for 20 min. Rate constants for FRET-dependent fluorescence were extracted using Origin software by fitting the curves to a one-phase association exponential function:

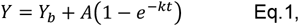

where Y_b_ is baseline, A is the amplitude, *k* is the rate constant, and t is time. At least two independent tests were carried out for each experiment.

### ATPase assay

SecA ATPase activity assay was done using malachite green kit (MAK307, Merck/Sigma). Briefly, 25 nM or 0.5 μM SecA was mixed with 0.5 μM SecB, 1 μM proOmpA, 10 mM DTT, 5 mM MgCl_2_ in presence of 50 mM Hepes/KOH pH 7.4 and 150 mM KCl. 0.125 or 1 mM SecYEG proteoliposomes (or 1 μM SecYEG) were added, the reaction was started by the addition of ATP, and incubated for 15 or 30 minutes at 37 °C. The reaction was diluted 10 times when necessary. Afterwards, the reaction was stopped by the addition of 20 μL working reagent supplied with the kit. The color was allowed to develop for 30 minutes at the ambient temperature, and the absorbance was measured at 590 nm. The data were fitted to Michaelis-Menten equation:

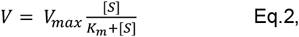

where V is the reaction velocity, V_max_ is the maximum reaction velocity, [S] is the ATP concentration, and Km is the Michaelis’s constant. At least two independent tests were carried out for each experiment.

### SecA flotation assay

SecA was mixed with liposomes in protein:lipid molar ratio of 1:5000, and the volume was completed to 100 μL with the buffer (50 mM KCl, 50 mM Hepes/KOH pH 7.4, 5 mM MgCl_2_). When mentioned, elevated concentrations of KCl were used. Samples were incubated 30 minutes at 25 °C and then mixed with 60 % sucrose (w/v) to achieve final sucrose concentration of 30 %. The samples were loaded in the centrifugation tube (S12-AT3 rotor, Thermo Fisher/Sorvall), followed by 20 % sucrose solution (250 μL) and the sucrose-free buffer (50 μL). Samples were centrifuged for 1 h at 80,000 rpm (AT3 rotor). Samples were then fractionated into 3 fractions, top (125 μL), middle (125 μL) and bottom (250 μL), then precipitated by the addition of 200 μL of 20 % TCA. The pellets were resuspended in the sample buffer and loaded on SDS-PAGE. The intensity of Coomassie-stained bands was quantified (ImageQuant TL, Cytiva/GE Life Sciences), and the amount of SecA bound to liposomes was determined by dividing the intensity of the floating fraction (top) by the integral intensity of all fractions. At least two independent tests were carried out for each experiment.

### Surface plasmon resonance

SecA:lipid binding experiments were performed using L1 sensor chip on two-channel Biacore X100 instrument (GE Healthcare Life Sciences). DOPC-only liposomes lacking SecA interaction were employed in the reference channel. SPR relies on changes in the evanescence wave within a short distance, typically ~100 nm, above the sensor surface, so the liposomes were additionally pre-extruded to the diameter of 50 nm. Prior the experiment, the chip surface was cleaned with 20 mM CHAPS (2 injections for 30 s each) and conditioned using SPR running buffer (150 mM KCl, 5 mM MgCl_2_, 50 mM Hepes/KOH pH 7,4). Liposomes were immobilized at the flow rate of 5 μL/min for 600 s to achieve 7,000 to 10,000 response units. Afterwards, 2 injections of 100 mM NaOH were performed to remove loosely attached material. SecA was transferred into the SPR running buffer and was injected for 150 s at the flow rate of 10 μL/min. When indicated, 300 mM KCl was used instead of 150 mM KCl in SPR running buffer to probe SecA binding at higher ionic strength. After each measurement, the chip surface was regenerated (2 injections of 20 mM CHAPS), so the immobilized liposomes were removed. Data were fitted by non-linear regression analysis of response levels at the steady state to one-site-binding model:

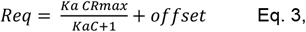

where *K*a is the association constant, C is the concentration of SecA, R_max_ is the maximum response unit, and *offset* is the intercept of the fitted curve on the y-axis. Due to extensive binding of SecA to liposome surfaces and potential dissociation/re-binding events known as mass transfer effect, a detailed analysis on binding/dissociation kinetics was omitted.

### Quartz crystal microbalance

Quartz crystal microbalance (QCM) measurements were carried using Q-Sense Omega Auto instrument (Biolin Scientific). This technique allows the real-time monitoring of SecA interactions with planar supported lipid bilayers (SLBs) by measuring the shifts in the resonance frequency and dissipation energy, which proportionally depend on the mass changes and changes in viscoelastic properties on the surface of the chip, respectively. The formation of SLBs on the QCM-D sensor chip (QSX 303 SiO2) was performed as follows. The surface of the plasma-treated sensor chip was equilibrated with aqueous buffer (50 mM Tris/HCL pH 7.4, 150 mM KCl) for 3 minutes to stabilize the frequency and dissipation energy baselines. Freshly extruded liposome suspension (extrusion via 100 nm membrane) was injected over the chip surface at flow rate of 50 μL/min for 5 min in 50 mM Tris/HCl pH 7.4, 150 mM KCl, 10 mM CaCl_2_. The liposomes could adsorb on the chip surface and underwent spontaneous collapse, which resulted in formation of SLBs. To remove loosely bound material from the chip surface, the surface was subsequently washed with the buffer (50 mM Tris/HCL pH 7.4 and 150 mM KCl). SLB formation resulted in a frequency shift of −27 ± 1 Hz and the energy dissipation of 0.7 ± 0.1, which is in excellent agreement with previously published data [49]. To probe SecA interaction with the SLBs, the ATPase was transferred into the running buffer (50 mM Tris/HCL pH 7.4, 150 mM KCl, 5 mM MgCl_2_) and the protein solution was then injected over the SLBs at flow rate of 50 μL/min. To determine the affinity of SecA to lipids, the concentration of the ATPase was varied in dilution series from 40 nM to 5.25 μM concentration, and the maximum change in the chip oscillation frequency was measured. Prior each cycle, the surface of the SLB was washed with the high-salt buffer (1 M NaCl, 20 mM Tris/HCL pH 7.4) and then with the running buffer to remove the bound SecA and equilibrate the SLB surface for the next round. The equilibrium dissociation constant (K_d_) for this measurement was determined by plotting steady-state net-frequency signal responses prior to dissociation phase (R_max_) against the corresponding SecA concentration (*C*). Data were fitted using GraphPad Prism 9 based on one-site binding model:

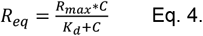

### Membrane fluidity analysis

Liposomes were mixed with 0.1 μM 1,6-diphenyl-1,3,5-hexatrien (DPH) to achieve dye:lipid ratio of 1:1000. Samples were incubated for 1 h at 25 °C in the dark. DPH fluorescence was recorded at 428 nm (slit width 5 nm) with the excitation wavelength of 350 nm (slit width 5 nm) using Fluorolog-3 fluorometer. Steady state anisotropy (r) was calculated as:

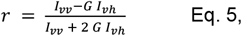

where *I* is the fluorescence intensity, v and h denote the vertical and horizontal setting for the excitation and emission polarizers, respectively, G is the instrumental correction factor which is provided by the instrument for each measurement. At least two independent tests were carried out for each experiment.

### Lipid packing analysis

Liposomes (lipid concentration 100 μM) were mixed with 0.3 μM laurdan to achieve dye:lipid ratio of 1:333. Samples were incubated at 37 °C for 1 h in the dark in presence of 50 mM KCl, 5 mM MgCl_2_, 50 mM Hepes/KOH pH 7.4. Laurdan emission spectrum was recorded from 400 to 600 nm (slit width 3 nm) with the excitation wavelength of 350 nm (slit width 3 nm) using Fluorolog-3 fluorometer. Generalized polarization value (GP) was calculated as a ratio of integrated intensities from 400 nm to 460 nm (*l*_1_) and from 470 nm to 550 nm (*l*_2_):

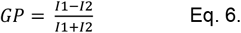

At least two independent tests were carried out for each experiment.

### Cardiolipin head group parametrization

The parameters for the cardiolipin head group were obtained following the linings established in Lipid11 [50] and Lipid14 [51]. Briefly, multiple conformations of the methyl-capped headgroup were generated with Balloon, using an RMSD cutoff of 1 Å. The resulting 21 independent structures were optimized, and the electrostatic potential (ESP) was computed, using Gaussian 09 at the HF/6-31G* theory level, with parameters as given by antechamber [52]. The resulting ESP for all conformations were combined into a multi-conformational fit, fixing the capping methyl group charges as established in Lipid11 [50] and using a standard two-step RESP procedure [52]. The obtained partial atomic charges were used together with Lipid17 atom types to generate an AMBER force field library file with LEaP. As the head group has four positions where acyl chains have to be attached (compared to the standard two attachment points per residue in AMBER), explicit bonds have to be set for two positions per cardiolipin when parametrizing a membrane system in LEaP. The parameters have been included in PACKMOL-Memgen [53] of AMBER20, where all combinations of the headgroup with every possible acyl chain in Lipid17 have been considered.

### MD simulations

Systems for molecular dynamics simulations were prepared with PACKMOL-Memgen [53], using a length of the membrane in x and y direction of 100 Å and default options otherwise. To mimic the experimental conditions, compositions of DOPC:DOPG 70:30, POPC:POPG 70:30, POPC:POPG:TOCL 70:25:5, POPC:POPG:bis(PO)CL 70:25:5, POPC:bis(PO)CL 85:15 and POPC:TOCL 85:15 were prepared. In addition, systems of DOPC:DOPG 70:30 and POPC:POPG 70:30 including the 25 N-terminal residues of SecA with an N-methyl amide cap in the C-terminus at 25 Å of the membrane surface were prepared. The peptide structure was modelled with TopModel [54], using as main templates PDB IDs 3BXZ (*E. coli* SecA) and 3DL8 *(B. subtilis* SecA). In all cases, potassium ions were added to neutralize the charges introduced by the negatively charged headgroups. To ensure independent starting configurations, all systems were packed five times with a different random seed.

From the packed systems, independent replicas were energy-minimized using the pmemd implementation included in AMBER18 [55], using ff14SB [56], TIP3P [57], and Lipid17 [51,58] parameters for the protein, water, and membrane lipids, respectively. To relax the system stepwise, alternating steepest descent/conjugate gradient energy minimizations with a maximum of 20,000 steps each were performed. Initially, the positions of the membrane were restrained during minimization; the final round of minimization was performed without restraints. To thermalize the systems, a Langevin thermostat [59] with a friction coefficient of 1 ps^-1^ was used. The pressure, when required, was maintained using a semi-isotropic Berendsen barostat [60] with a relaxation time of 1 ps, coupling the membrane (x-y) plane. The system was heated by gradually increasing the temperature from 10 to 100 K for 5 ps under NVT conditions, and from 100 to 300 K for 115 ps under NPT conditions at 1 bar. The thermalization process was continued for 5 ns under NPT conditions, after which production runs of 1 μs length were performed using the same conditions with the pmemd GPU implementation [61], constraining covalent bonds to hydrogens with the SHAKE algorithm [62] and using a time step of 2 fs.

The trajectories were analysed with cpptraj [63] as to lipid order parameters and electron density profiles. The lipid order parameter describes the level of order imposed on the lipid molecules in a bilayer arrangement and relates with deuterium-NMR measurements [64]. The electron density profile describes the probability of finding electron-rich regions along the membrane normal and can be related to X-ray scattering experiments [65]. As the membrane has an anisotropic, i.e., planar arrangement, it gives information regarding the bilayer arrangement, which in simulations can be additionally decomposed according to the contribution of each system component, obtaining information of its location along the membrane normal. In all cases, the profiles describe well-behaved membrane bilayers (Suppl. Figure 3). The contacts of the SecA N-terminal peptide with the membrane headgroups were evaluated as the sum of the per-residue contributions as obtained from the native contacts routine, using a cut-off of 4.5 Å. The average area per lipid of each system was calculate with cpptraj from the area of the xy plane and the number of lipids on each leaflet. To measure the per-lipid type contribution to the area per lipid, the APL@Voro software was used [66]. For this, the trajectories were centered and imaged on the bilayer, and transformed into the GROMACS XTC format with cpptraj. Afterwards, the trajectories were processed with the software, assigning the phosphorous atoms (or the central carbon of the glycerol moiety of cardiolipins) to the area by tessellation. The average xz particle density was calculated using the volmap function of cpptraj, which represents each atom as a gaussian with a standard deviation equal to the atomic radius, similarly to what has been described previously [13]. Briefly, a 80×80×80 Å grid centered in the membrane center, with a 1 Å spacing in every dimension, and the average density was calculated along the 1 μs simulation of each system. To obtain the xz profile, the density along the y dimension was averaged.

To calculate the lateral pressure of the equilibrated systems after 1 μs of simulation time as a function of the z-coordinate, all replicas were extended for additional 100 ns, recording the coordinates and velocities every 5 ps. The obtained trajectory was centered on the bilayer, transformed to GROMACS TRR format with cpptraj, and post-processed with GROMACS-LS to obtain the stress tensors [67]. For this, a non-bonded cut-off of 20 Å and otherwise equivalent conditions to the production run were used, processing each ns of simulation independently, and calculating the average stress tensor with the provided tensortools script. The lateral pressure was calculated according to equation E.5

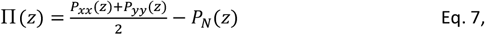

where the first term corresponds to the average lateral term (*P*_L_), and *P*_N_ corresponds to the normal component.

## Acknowledgements

We thank Alexander Büll and Nicola Vettore (DTU Denmark) for the assistance with the QCM experiments. The research was supported by the Deutsche Forschungsgemeinschaft (DFG) via the research grant KE1879/3-1 to A.K. and projects A10 (A.K.) and A03 (H.G.) within the CRC 1208 “Identity and dynamics of biological membranes” (project number 267205415). We are grateful for computational support and infrastructure provided by the “Zentrum für Informations-und Medientechnologie” (ZIM) at the Heinrich Heine University Düsseldorf and the computing time provided by the John von Neumann Institute for Computing (NIC) to H.G. on the supercomputer JURECA at Jülich Supercomputing Centre (JSC, user IDs: plaf and HKF7).

## Author contributions

MK performed biochemical and fluorescence-based analysis; MK and ML carried out SPR and DSF measurements; ML carried out QCM experiments; MK, ML and AK analyzed and interpreted the data; SSV and HG carried out the computational simulations and interpreted the data in cooperation with MK and AK; all authors contributed to writing and editing the manuscript.

## Conflict of interest

Authors declare no conflict of interest in relation to the presented work.

## Supplementary figures

**Supplemental Figure 1.**
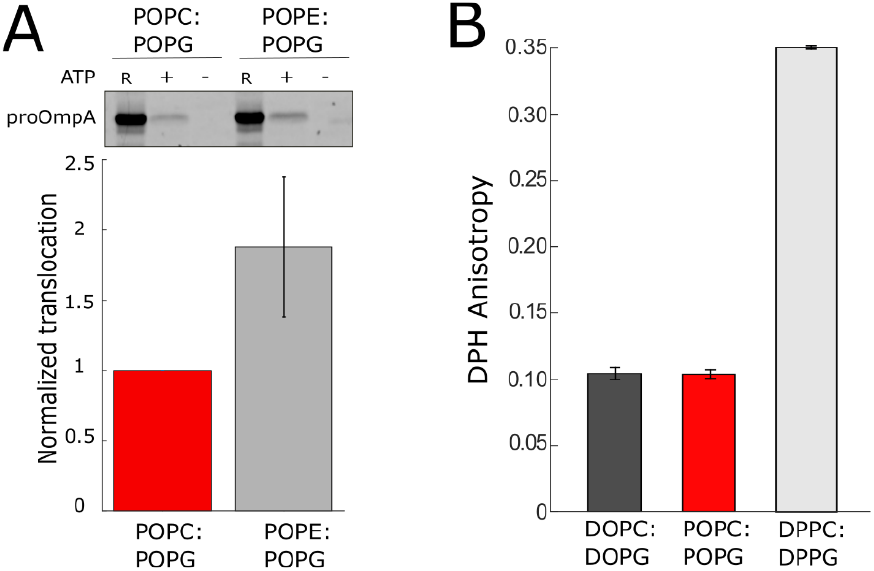
**(A)** SecYEG remains active in POPC:POPG liposomes. Translocation of the model substrate proOmpA was measured in SecYEG-containing proteoliposomes, which contained either 70 mol % POPC or POPE zwitterionic lipids. **(B)** Both DOPC:DOPG and POPC:POPG lipid bilayers are present in the fluid phase at 37°C, as probed by DPH fluorescence anisotropy. DPPC:DPPG membranes form low-mobile gel phase at these conditions, which results in a high fluorescence anisotropy value.

**Supplemental Figure 2.**
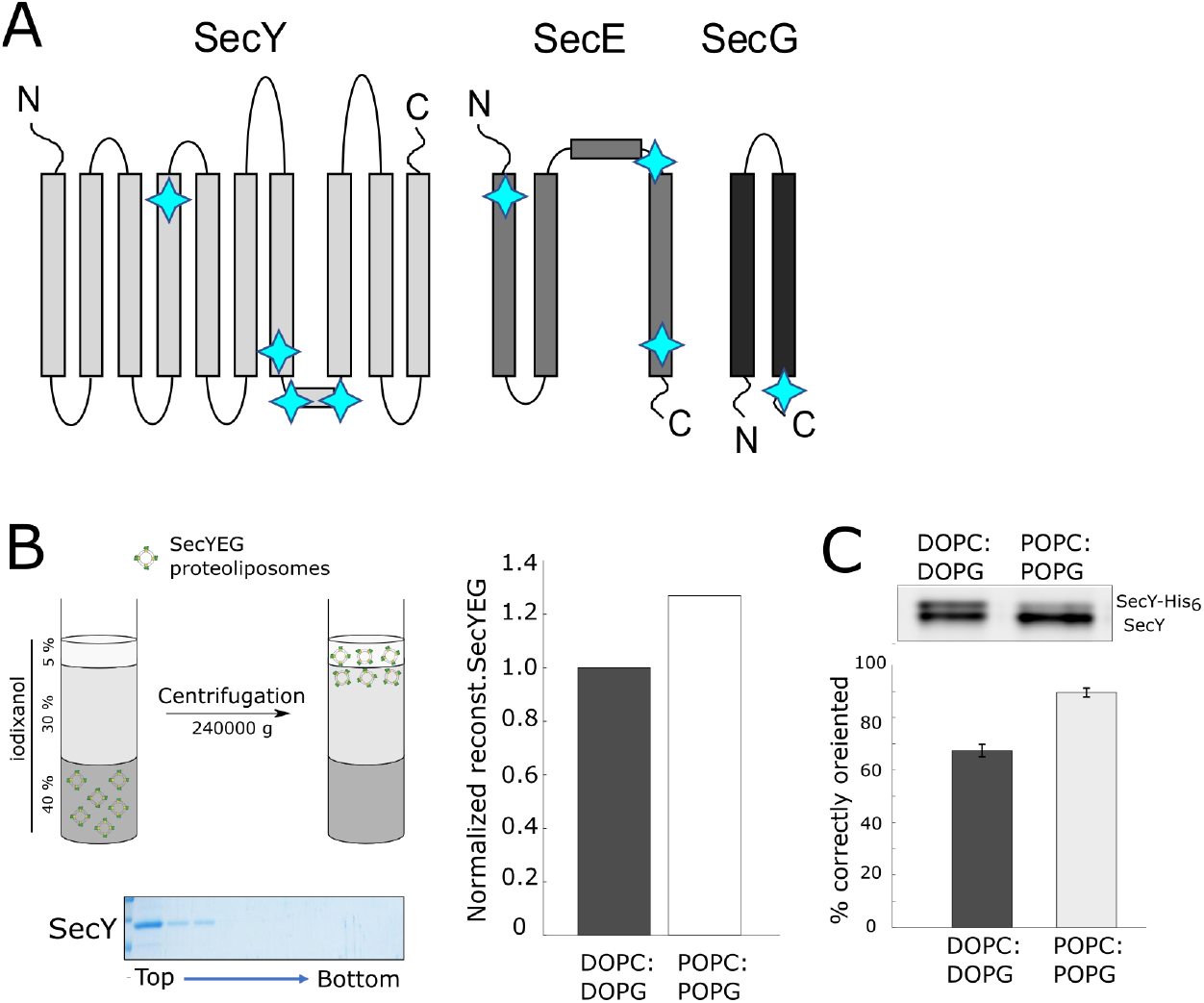
Analysis of SecYEG folding and stability in proteoliposomes. **(A)** The scheme of the SecYEG secondary structure. Positions of the tryptophan residues are indicated with star symbols. Changes of tryptophan fluorescence were used to describe the thermal unfolding of SecYEG in DSF experiments. **(B)** Flotation assay in iodixanol density gradient using SecYEG proteoliposomes demonstrated nearly equal reconstitution efficiency of the translocon for DOPC:DOPG or POPC:POPG lipids. The example SDS-PAGE shows the distribution of SecY reconstituted into DOPC:DOPG liposomes. **(C)** Site-specific cleavage of the N-terminal poly-histidine tag of SecY subunit by enterokinase reports on the accessibility of the tag in liposomes, and so reveals the orientation of reconstituted SecYEG. In POPC:POPG liposomes nearly 80 % of the translocons were exposed to the protease, and so acquired the functionally-relevant orientation.

**Supplemental Figure 3.**
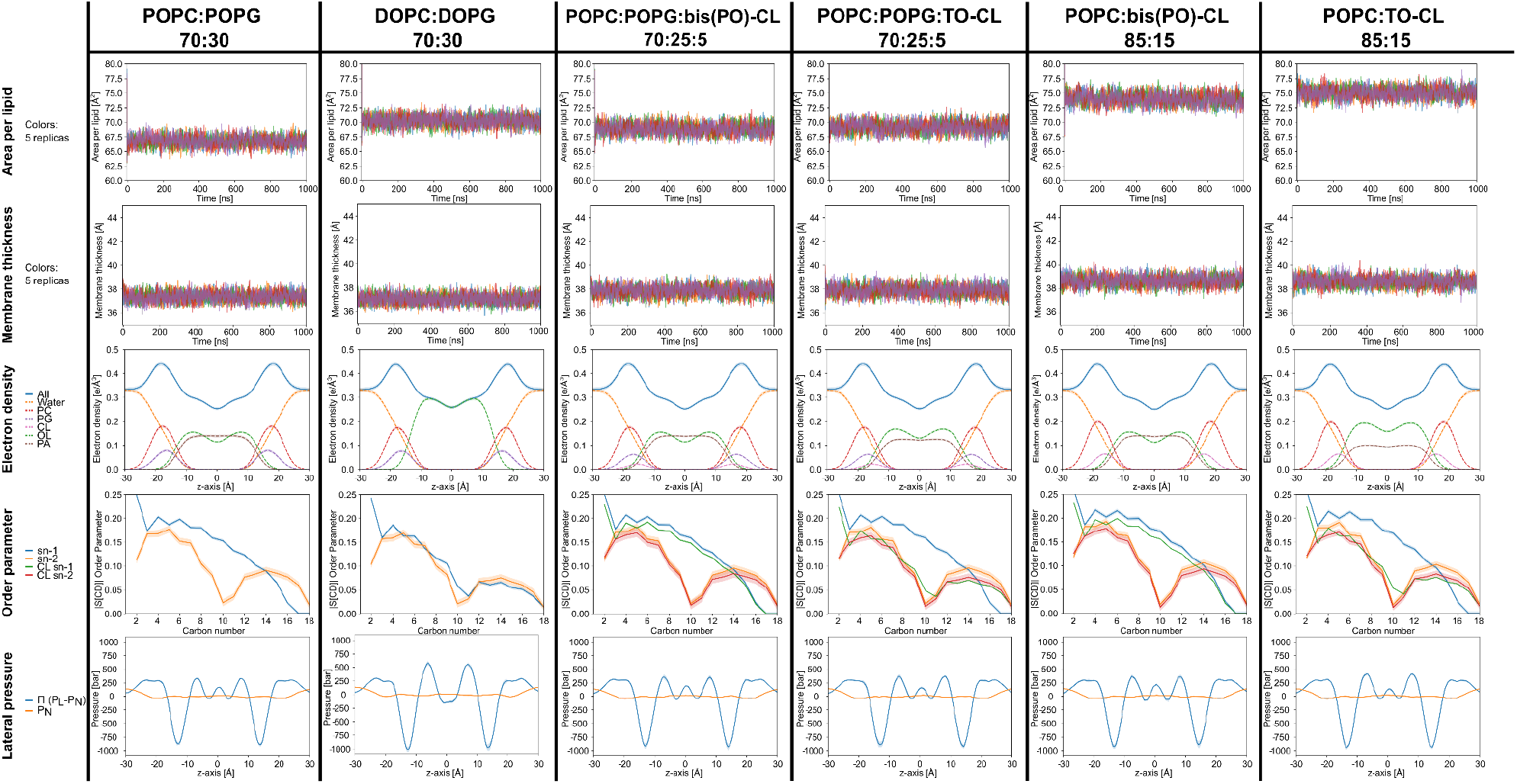
Biophysical properties of lipid bilayers determined from all-atom MD simulations. Area per lipid, membrane thickness, electron density profiles, order parameter, and lateral pressure profiles obtained by MD simulations for all investigated lipid bilayers. Shaded areas represent the standard error of the mean (SEM). Mean values ± SEM of the area per lipid and thickness of the bilayers can be found in Table 1.

**Supplemental Figure 4.**
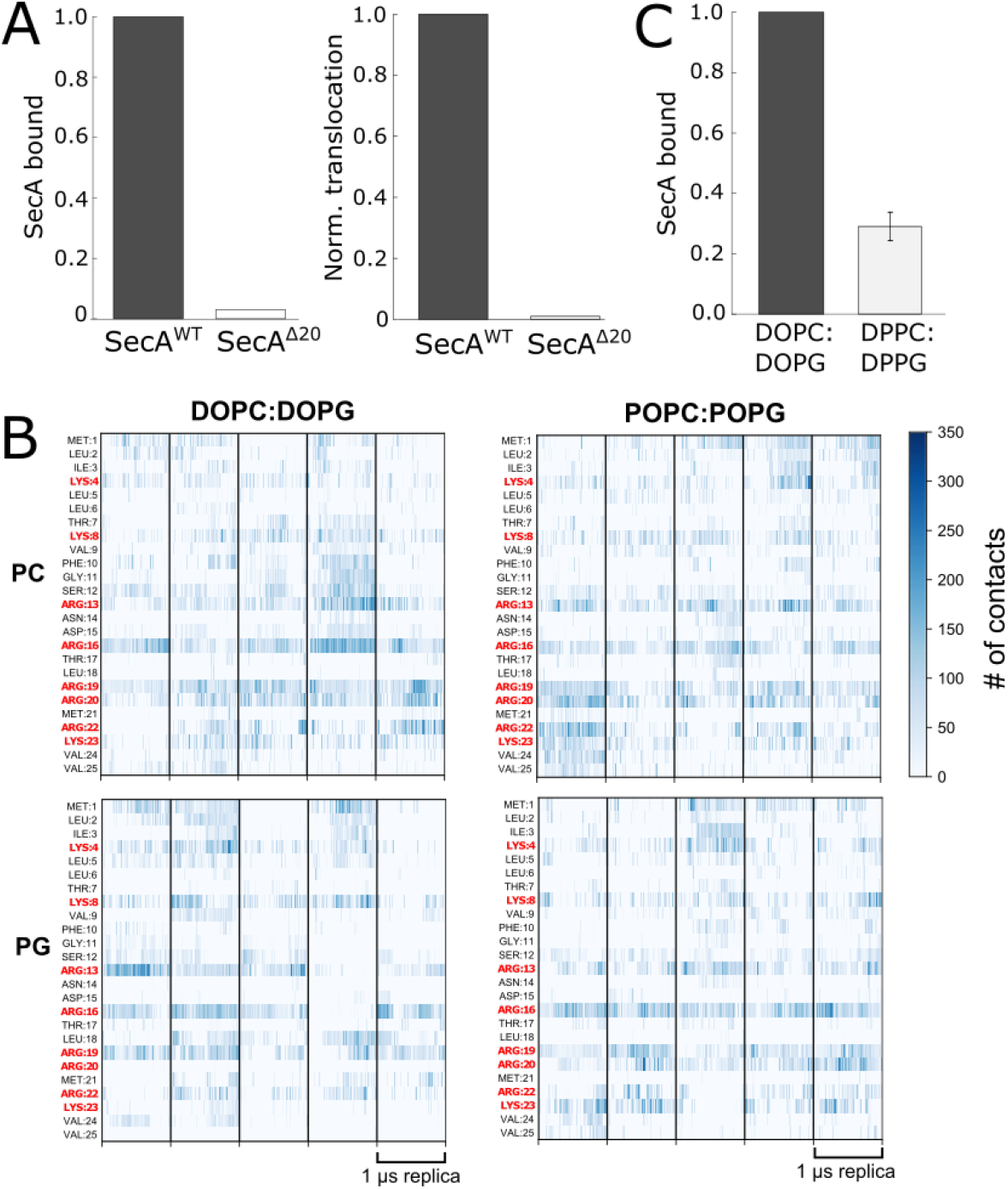
SecA:lipid interactions are determined by the N-terminal domain. **(A)** Deletion of SecA N-terminal helix abolishes lipid binding as tested via the flotation assay and eliminates SecA:SecYEG translocation activity in DOPC:DOPG liposomes. **(B)** Interactions of the SecA N-terminal helix (residues 1 to 25) with DOPC:DOPG and POPC:POPG membranes over MD simulations of 1 μs length. Per-residue contacts with membrane headgroups in each replica are shown. Cationic residues are highlighted in red. **(C)** Flotation assay shows that SecA binding to gel phase liposomes composed of DPPC:DPPG lipids is strongly suppressed in comparison to fluid phase liposomes (DOPC:DOPG), despite the identical content of anionic lipids.

**Supplemental Figure 5.**
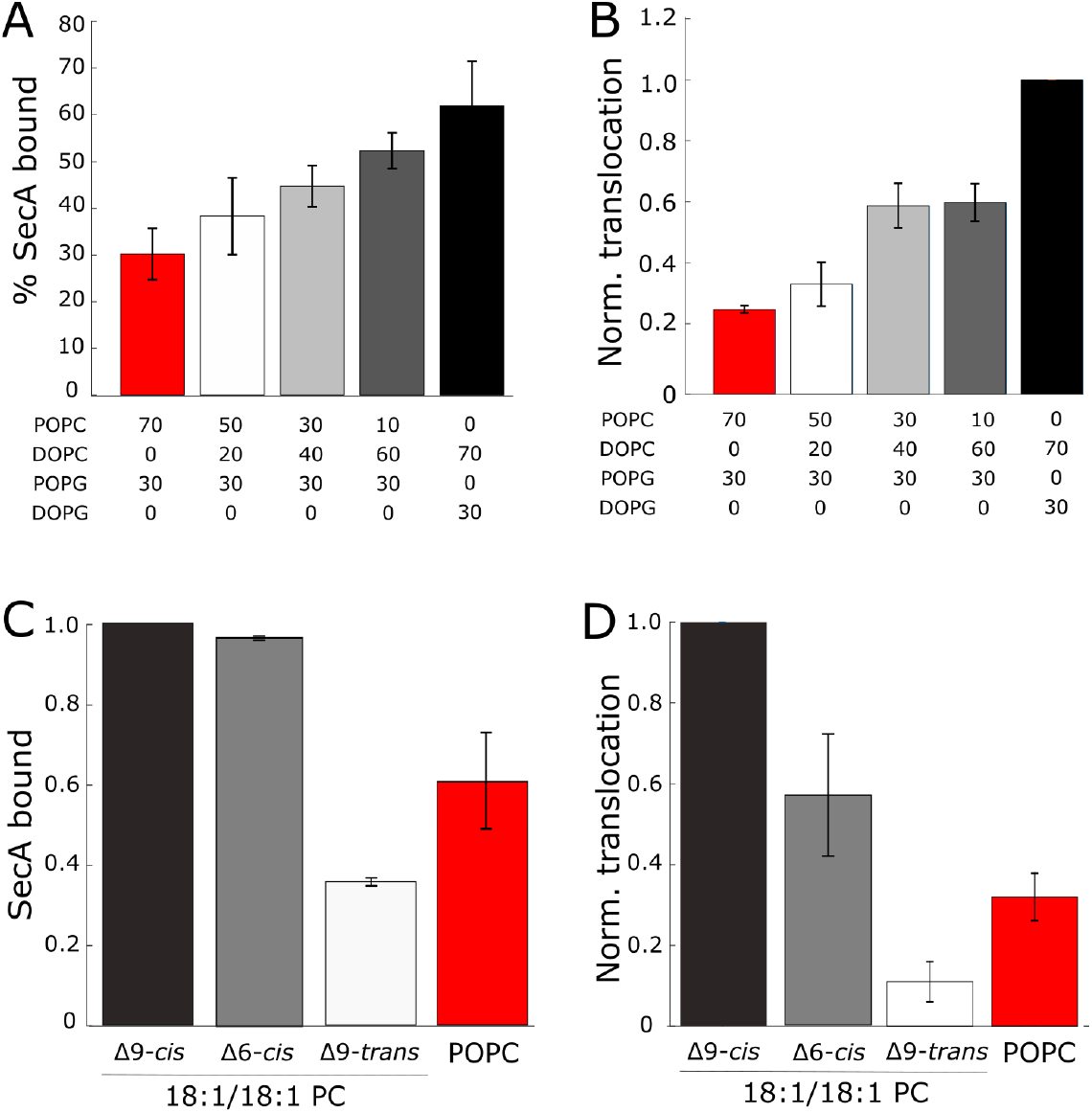
Changes in SecA:lipid binding upon alternations in the UFA content correlate with SecA:SecYEG activity. **(A)** Upon gradual increase in the UFA content the SecA:lipid binding (flotation assay) correlates with the translocation activity of SecA:SecYEG complex (**B**, protease protection assay). **(C)** SecA binding is stimulated by *cis-*UFAs and is hindered in the presence of Δ9-*trans* UFA. All liposomes contained 30 mol % POPG and 70 mol % indicated PC lipids. **(D)** The translocation activity of the SecA:SecYEG complex in different types of liposomes correlates with SecA binding to the liposomes (Suppl. Figure 5C) and the lipid packing order, measured via laurdan fluorescence polarization (Figure 3A). Thus, the configuration of the unsaturated fatty acid determines the functionality of SecA:SecYEG complex.

**Supplemental Figure 6.**
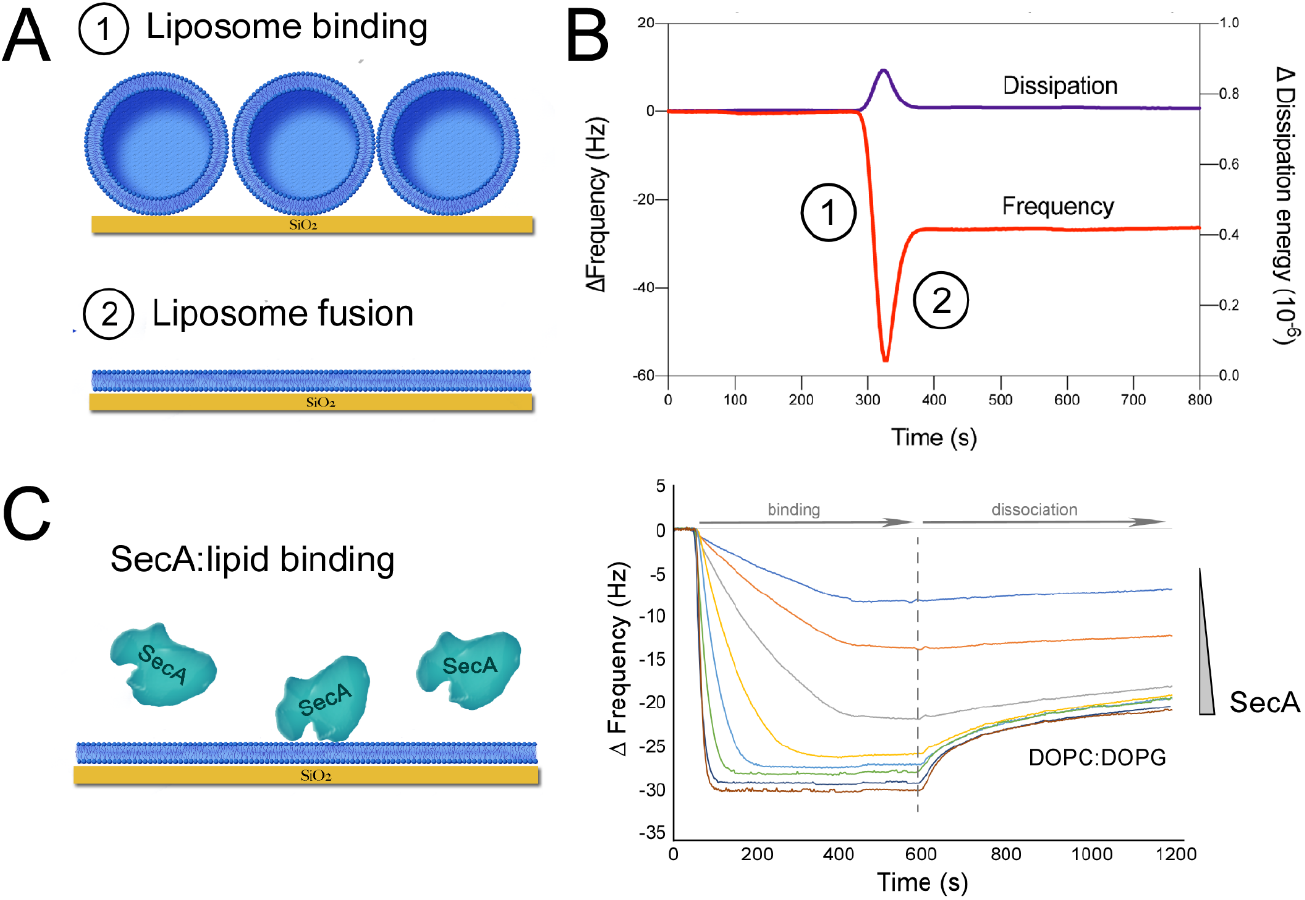
Quartz crystal microbalance to probe SecA:lipid interactions. **(A)** Formation of the lipid bilayer on the QCM chip surface. Liposomes are bound to the surface at high density, and their fusion in the presence of calcium leads to the formation of the supported lipid bilayer. **(B)** The attachment and the fusion of liposomes are observed as changes in the oscillation frequency of the chip (stages 1 and 2). An increase in the energy dissipation upon the liposome attachment is due to the large volume of encapsulated water, which is further released upon fusion (stage 2). **(C)** Measuring SecA binding to the lipid bilayer via changes in the oscillation frequency. SecA was injected in the buffer flow over DOPC:DOPG bilayer to monitor the association and dissociation stages. SecA concentration ranged from 40 nM to 5.25 μM (two-fold dilution per titration step).

**Supplemental Figure 7.**
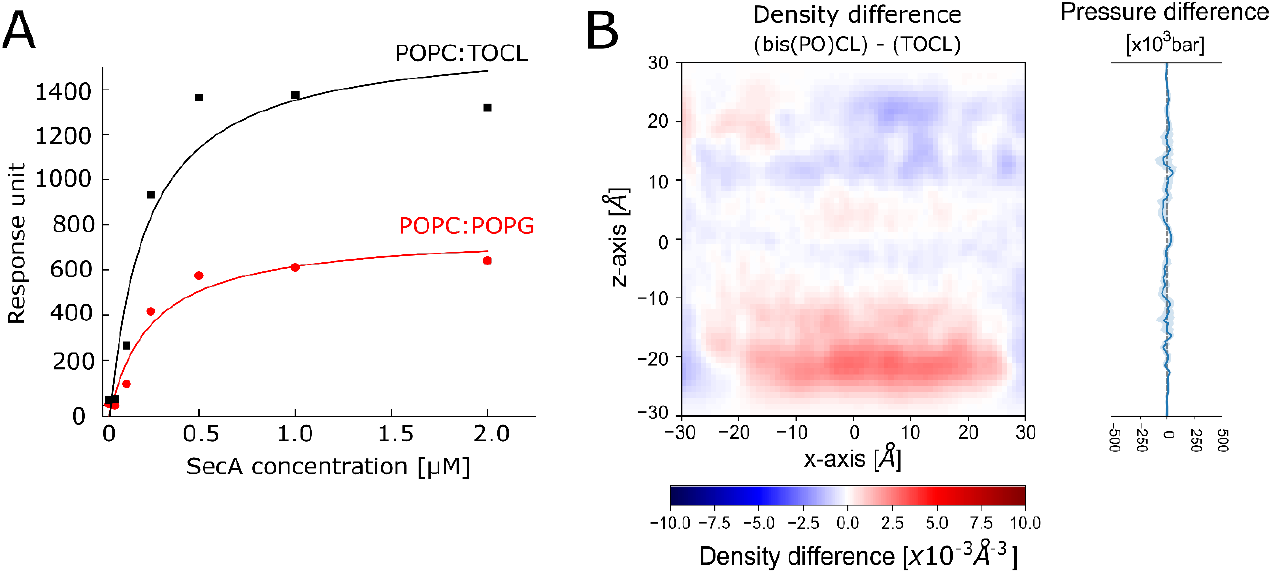
**(A)** SPR experiments show enhanced binding of SecA to liposomes containing 15 mol % TOCL in comparison to liposomes with 30 mol % POPG, despite the identical surface charge. **(B)** 2D lipid density (left) and lateral pressure (right) differences obtained by MD simulations between bilayers composed of POPC:POPG:bis(PO)CL and POPC:POPG:TOCL (molar ratio 70:25:5) reveal only minor differences compared to those observed at higher concentrations of CL (Figure 6D) or when changing the UFA composition of the whole bilayer (Figure 3E).

